# Complete Genomes of Cultivated Gut Bacteria Reveal Mobile Genetic Element-Driven Functional Diversity with Therapeutic Implications

**DOI:** 10.64898/2026.03.25.714112

**Authors:** Haoyu Wang, Yuzheng Gu, Wenxin He, Jinlong Yang, Hewei Liang, Mengmeng Wang, Zhinan Wu, Yangfeng Wen, Jinhong Wang, Xing Rao, Yaqi Fan, Jiayi Ma, Xinyu Yang, Xin Tong, Liye Yang, Yu-sheng Xu, Juan Zhao, Tao Zeng, Yuning Zhang, Yiyi Zhong, Haifeng Zhang, Chuan Liu, Xuechun Shen, Karsten Kristiansen, Juan Yang, Fushu Liu, Zejun Yang, Wangsheng Li, Peng Gao, Jun Xu, Shangyong Li, Ningning He, Bo Wang, Xin Jin, Chuanyu Liu, Xun Xu, Yuliang Dong, Hongwei Zhou, Feng Zhao, Liang Xiao, Yuanqiang Zou

**Author notes:** These authors contributed equally to this work (Haoyu Wang, Yuzheng Gu, Wenxin He, Jinlong Yang). Correspondence: Dr. Yuanqiang Zou Dr. Liang Xiao Dr. Feng Zhao Dr. Hongwei Zhou Dr. Yuliang Dong.

## Abstract

Cultivated bacterial isolates are essential for elucidating gut microbiota functions, yet reliance on fragmented draft genomes and limited mobile genetic element (MGE) annotation has constrained high-resolution analyses. Here, we present the Cultivated Complete Genome Reference (CCGR), a comprehensive compendium of 1,150 fully circularized human gut bacterial genomes. Overcoming draft limitations, CCGR achieves chromosome-scale resolution and reveals distinct evolutionary strategies shaping gene topological organization in response to bacterial growth. Furthermore, we decode the dynamic landscape of MGEs, demonstrating that broad-host-range phages follow strict positional patterns by integrating at conserved chromosomal hotspots flanked by nutrient transporter loci. Importantly, we provide mechanistic in vivo evidence that accessory genomic elements drive strain-level functional heterogeneity. Specifically, the plasmid-encoded scrK gene in Levilactobacillus brevis involved in fructose catabolism is required to mitigate fructose-diet-exacerbated colitis. Collectively, CCGR establishes a framework linking chromosomal architecture and MGEs to bacterial evolution and host health, enabling precise, strain-resolved functional investigations.

## Introduction

The intricate composition of the gut microbiota and its metabolic activities are fundamental to human health^1,2^. Genomic characterization of gut bacteria serves as a cornerstone for decoding their functional potential, which is important to understand the relationship between host and bacteria^3,4^. Studies of bacterial genomes can reveal strain-level genetic relationships such as phylogenetic diversity, distribution across different cohorts, horizontal gene transfer (HGT) dynamics, and the landscape of mobile genetic elements (MGEs)^5^.

In recent years, significant advancements in gut microbial genomic databases have expanded our understanding of microbiota-host interactions, with comprehensive resources collectively contributing to the characterization of bacterial diversity, functional gene landscapes, and metabolic potential^6–14^. Metagenomic assembled genomes (MAGs) and culturable draft genomes have enabled analyses of the relationship between human health and many microorganisms, and how probiotics may exert beneficial effects on human health^15–17^. Detailed analyses of microbial pan-genomes have also provided in-depth information on the relationship between microbial evolution and environmental adaptation^18–20^. These studies have not only enriched the catalogs of bacterial genomes but also facilitated research into carbohydrate-active enzymes, secondary metabolites, and probiotic candidates through integrated approaches combining metagenomic assembly, isolate pairing, and preclinical validation.

However, reliance on short-read sequencing can lead to fragmentation in the generated draft genomes and the median contig N50 is less than 500 kbp in major databases, which limits analyses of chromosomal organization and MGEs^9^. Furthermore, short-read–based draft genomes are often insufficient for resolving repetitive regions in bacterial chromosomes and preserving accurate positional information^21^. For example, terminal inverted repeats (TIRs) that encode biosynthetic gene clusters (BGCs) are frequently misassembled or absent in draft genomes, which can substantially compromise functional analyses^22^. Moreover, the chromosomal location of functional genes can influence their expression, highlighting the importance of positional effects on bacterial gene expression profiles^23^. In addition, genome fragmentation resulting from incomplete assemblies often impedes the accurate classification of extrachromosomal elements, such as plasmids and phages, and can bias diversity analyses^24,25^.

While recent advances in long-read metagenomics have significantly enhanced our resolution of the gut microbiome, this approach cannot yield viable isolates for downstream functional validation^26–29^. Furthermore, metagenome-assembled genomes (MAGs) remain susceptible to misassemblies, highlighting the continued necessity of complete, closed genomes derived from cultured single strains^30^. Therefore, the construction of complete bacterial genome references is essential for comprehensive and accurate studies of microbial ecology.

Here, we present the Cultivated Complete Genome Reference (CCGR), a curated resource of 1,150 circularized human gut bacterial genomes. Overcoming the limitations of fragmented draft assemblies, this chromosome-scale resolution allows us to systematically investigate the topological principles governing functional gene distribution, revealing that divergent evolutionary strategies dictate genomic architecture in response to rapid growth. Furthermore, we decode the dynamic landscape of mobile genetic elements (MGEs) to show that broad-host-range phages integrate following strict positional determinism at conserved chromosomal hotspots. We also provide mechanistic in vivo evidence that the accessory genome drives strain-level functional heterogeneity, demonstrating that a plasmid-encoded scrK gene in *Levilactobacillus brevis* is essential for alleviating dextran sulfate sodium (DSS) and high-fructose-diet-induced colitis. Finally, we identified specific phages and plasmids as robust, novel biomarkers across diverse clinical cohorts. We envisage that this comprehensive reference will expand current capabilities in genomic analysis, refining insights derived from draft datasets to provide a powerful framework for future strain-level investigations into MGE-host interactions and gut microbial ecology.

## Results

### A cultivated human gut bacterial genome repository of 1,150 circular genomes

To construct a large-scale Cultivated Complete Genome Reference (CCGR), 1,093 bacterial isolates were cultivated. Then, the draft genome of each strain was assembled from short-read sequencing data, and the complete genome was assembled using a hybrid approach combining both short-read and long-read data^31^. After genome quality assessment, 966 complete genomes were obtained and 127 multi-isolated strains were reassembled into 184 circular metagenome assembled genomes (MAGs). Thus, a total of 1,150 circular genomes were used to construct the CCGR (Fig. 1a, Supplementary Data 1). We subsequently clustered the 1,150 genomes into 199 species-level clusters based on 95% average nucleotide identity (ANI). Notably, Bacillota_A represented most of the clusters (146 genomes, 58 clusters), and one cluster containing 3 genomes from the genus *Collinsella* could not be classified at the species-level, indicating previously unidentified species (Fig. 1a, Supplementary Data 2). To further evaluate the contribution of CCGR to bacterial complete genomes, we mapped the CCGR genomes against the human gut-derived 1,562 bacterial complete genomes from NCBI RefSeq database. Results showed that complete genomes from 127 bacterial species were underrepresented in NCBI RefSeq database (Fig. 1b), especially for species in three phyla Bacillota_A, Bacillota, and Bacteroidota (Supplementary Fig. 1a). Moreover, CCGR includes a substantial number of high-quality genomes of *Bifidobacterium infantis*, *Bifidobacterium pseudocatenulatum*, *Escherichia coli*, *Phocaeicola dorei*, and *Bifidobacterium adolescentis* (Supplementary Fig. 1b), which can be used for strain-level comparative genomic analysis.

**Fig. 1.**
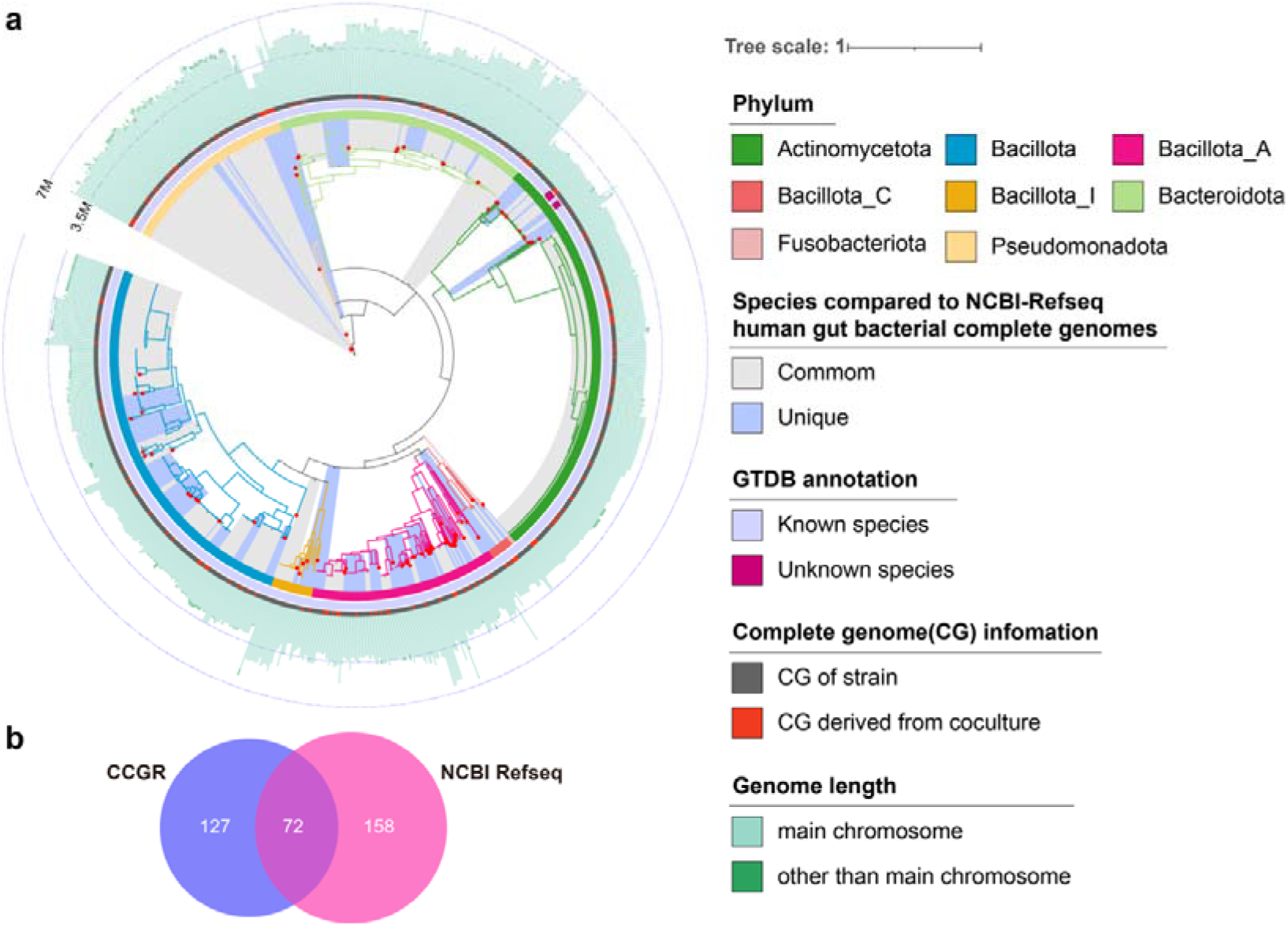
Taxonomic profile of CCGR. **a** Maximum likelihood phylogenetic tree of the 1,150 genomes of CCGR. Singleton genomes are marked with inner red dots. Color range indicates the common (light grey) and unique (light purple) species compared to NCBI-Refseq database. Tree branches and the first layer depict the GTDB phylum annotation, the second layer describes the matching to the GTDB database at species level, and the third layer shows the genome derived from pure culture (grey) and contaminated culture (red). The circumferential bar plot illustrates the genome size, whereas the main chromosome is shown in light green, and fragments other than main chromosome are shown in dark green. **b** Matching of CCGR2 to the NCBI-Refseq database at species-level. The Venn diagram is colored according to the origin of the species and the numbers are indicated.

### Complete genomes reveal divergent strategies in genomic architecture

Recent studies have established that bacterial genes exhibit a positional preference relative to the replication origin^32^. Using the complete genomes, we validated this topological constraint across the five major bacterial phyla in CCGR, confirming that the genomic distribution of highly prevalent genes is non-random, a trend particularly robust in the phylum Actinomycetota (Supplementary Fig. 2). Interestingly, Pseudomonadota genomes frequently exhibited symmetric inversions around the replication origin-terminus axis, a distinctive architectural feature that has been reported in a prior study^33^. Beyond this positional bias, we investigated strand-specific preferences of genes (Supplementary Fig. 3). Actinomycetota and Pseudomonadota exhibited a striking “U-shaped” bimodal profile, with genes strongly polarized to either the leading or lagging strand extremes. In contrast, Bacillota displayed a pervasive unidirectional skew towards the leading strand, whereas Bacteroidota showed a relatively uniform distribution, suggesting relaxed strand constraints (Supplementary Fig. 3).

To quantify the structural implications of these strand biases, we calculated the strand switch rate (the frequency of transcriptional orientation changes between adjacent genes) and revealed a marked structural dichotomy across phyla. Pseudomonadota exhibited the highest median strand switch rate, indicating a highly fragmented genomic architecture composed of short transcriptional units. In contrast, Bacillota_A and Bacillota maintained significantly lower switch rates, preserving longer co-directional gene arrays (Fig. 2a, b). This genomic fragmentation was density-dependent, meaning that in regions with a higher gene density, the strand switch rate was lower (Fig. 2c). Regions with the highest gene density harbored significantly longer arrays of co-directional genes compared to low-density regions, indicating that high gene density might impose a topological constraint to select for continuous transcriptional orientations (Fig. 2d). We further dissected these constraints across functional categories. Genes related to translation, ribosomal structure and biogenesis exhibited the most rigid architectural conservation, characterized by the highest leading strand bias and the lowest strand switch rate (Fig. 2e, f).

**Fig. 2.**
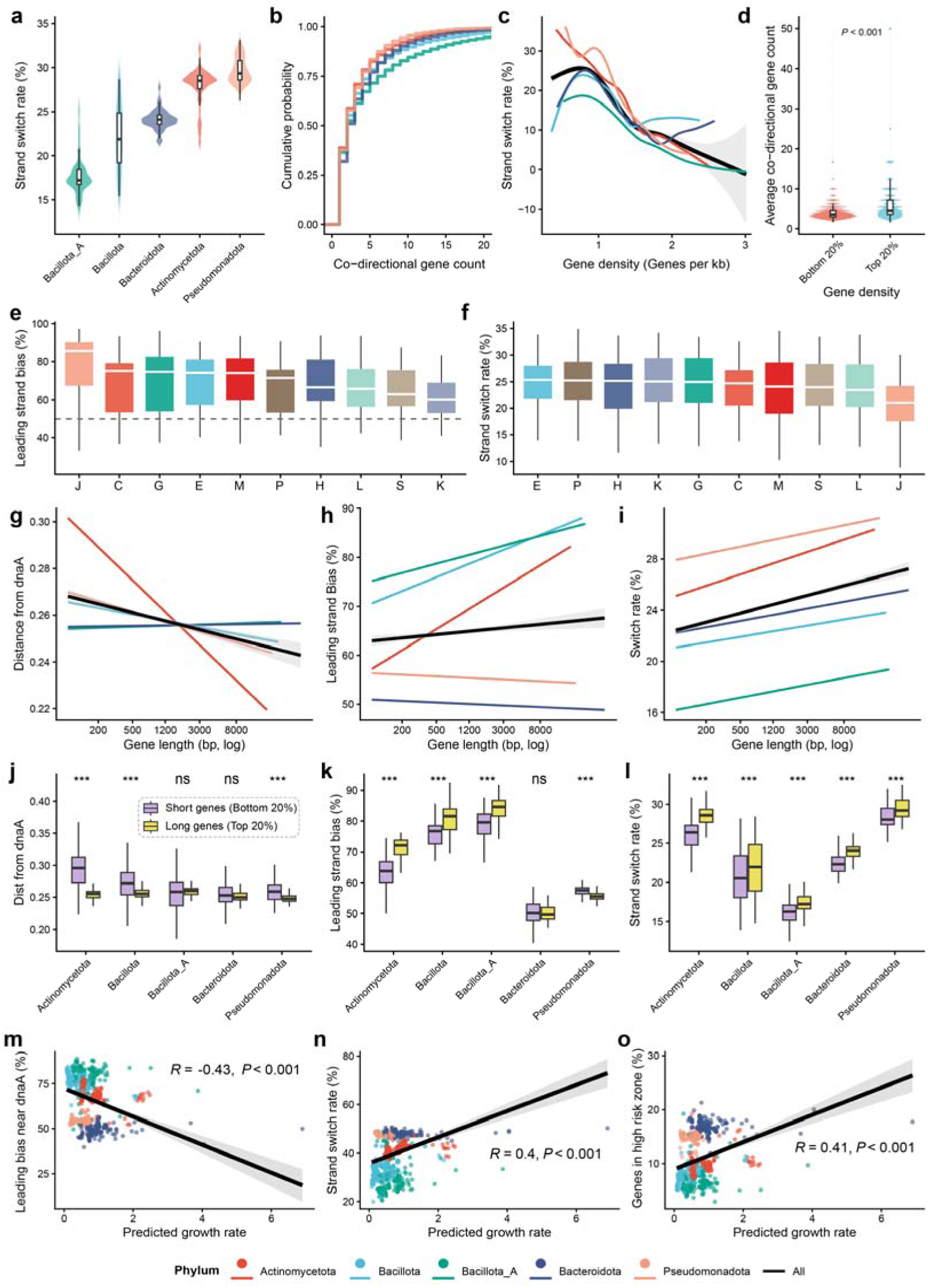
Complete genomes reveal divergent strategies in genomic architecture. **a** Distribution of genomic strand switch rates across different bacterial phyla. The strand switch rate is defined as the frequency of transcriptional orientation changes between adjacent genes. **b** Cumulative distribution function of co-directional gene counts across phyla, illustrating the probability distribution of operon-like gene clusters of varying lengths. **c** Relationship between local gene density (genes per kb) and strand switch rate. The trend lines represent LOESS regression fits with 95% confidence intervals (shaded areas). **d** Quantification of co-directional gene runs (consecutive genes on the same strand) in genomic regions with low (bottom 20%) versus high (top 20%) gene density. **e** Leading strand bias distribution across functional categories defined by Clusters of Orthologous Groups (COG). The dashed line at 50% indicates a random distribution between leading and lagging strands. **f** Strand switch rate distribution across COG functional categories. **g** Relationship between gene length and spatial distribution of genes relative to *dnaA*. The Y-axis represents the normalized distance from *dnaA*. Genes were binned by length (log-transformed). **h** Trend lines indicate the probability of a gene being located on the leading strand relative to its coding sequence length. **i** Dependence of strand switch rate on gene length. **j-l** Comparison of the normalized distance from *dnaA* (j), leading strand bias (k), and strand switch rate (l) between short (bottom 20%) and long (top 20%) genes across phyla. **m** Correlation between the predicted maximal growth rate and leading strand bias specifically within the replication origin-proximal region (defined as the genomic interval spanning the first one-third of the distance from *dnaA* to the replication terminus). **n** Correlation between predicted growth rate and the global strand switch rate, linking organismal growth dynamics to overall genomic fragmentation. **o** Correlation between predicted growth rate and the proportion of genes located in high risk zones. High risk zones are defined as genomic regions proximal to *dnaA* but located on the lagging strand, where replication-transcription conflicts are most severe. Asterisks indicate statistical significance (\*\*\**P* < 0.001, \*\**P* < 0.01, and \**P* < 0.05).

We next explored whether the observed genomic fragmentation is constrained by physical gene length. Globally, a conserved spatial bias where longer genes were preferentially enriched in the replication origin-proximal regions (Fig. 2g). As gene length increased, there were universal increases in leading strand bias and strand switch rate (Fig. 2h, i). However, the stringency of these constraints exhibited sharp phylogenetic divergence. Specifically, the spatial sequestration of long genes near the origin of replication was notably attenuated in Bacillota_A and Bacteroidota, where distance distributions for long and short genes were indistinguishable (Fig. 2j). Similarly, in Pseudomonadota and Bacteroidota, the topological confinement of long genes to the leading strand was effectively uncoupled (Fig. 2k). This topological sorting resulted in their lagging-to-leading gene length ratios being closed to unity, indicating that gene length is decoupled from strand specificity in these phyla (Supplementary Fig. 4). Despite these specific divergences, a unifying architectural feature across these phyla was that long genes consistently displayed higher strand switch rates compared to short genes (Fig. 2l).

Finally, to link genomic architecture with life history traits, we correlated architectural metrics with predicted maximal growth rates of genomes. Generally, rapid growth appeared to impose a heavy topological cost, correlating with a paradoxical reduction in leading strand bias near the origin (Fig. 2m), increased strand switch rates (Fig. 2n), and the accumulation of genes in regions proximal to *dnaA* but located on the lagging strand (Fig. 2o). However, phylogenetic decomposition revealed that this trade-off was driven predominantly by Bacillota and Actinomycetota (Supplementary Fig. 5a-c). In contrast, Pseudomonadota genomes defied these constraints, exhibiting a robust architectural framework where rapid growth was decoupled from topological constraints. Specifically, Pseudomonadota maintained stable origin-proximal bias and displayed a negative correlation between growth and switch rate, thereby preserving architectural order even under high replication pressure.

### Complete genomes reveal a refined genetic structure of gut bacteria

Harnessing long-read sequencing technology enables bacterial genomic analysis at a complete genome resolution. Here, to quantify the advancements in genomic integrity of complete genomes, we compared the counts of multiple genomic metrics between 966 complete genomes and paired draft genomes (Fig. 3a, Supplementary Data3). As expected, complete genomes harbored higher counts of bases, coding sequences (CDSs), and genes, exhibiting the most striking improvements in rRNA and tRNA gene counts. Specifically for each genome, the improvement of these parameters by the complete genomes was particularly evident in the genomes of the Bacillota, Bacillota_A, and Pseudomonadota phyla (Fig. 3d). We found that the increase in indicators such as the number of rRNA and tRNA genes was negatively correlated with the GC content and positively correlated with the genome size (Fig. 3e), suggesting that genome closure yields greater improvements in larger and low-GC genomes, which are prone to assembly gaps and gene loss in draft versions. Moreover, the complete genomes led to a marked enhancement in the quality of contig length and gene counts of phages and plasmids (Fig. 3b, c), suggesting complete genomes reveal more refined information regarding the interactions between mobile genetic elements (MGEs) and their host bacteria.

**Fig. 3.**
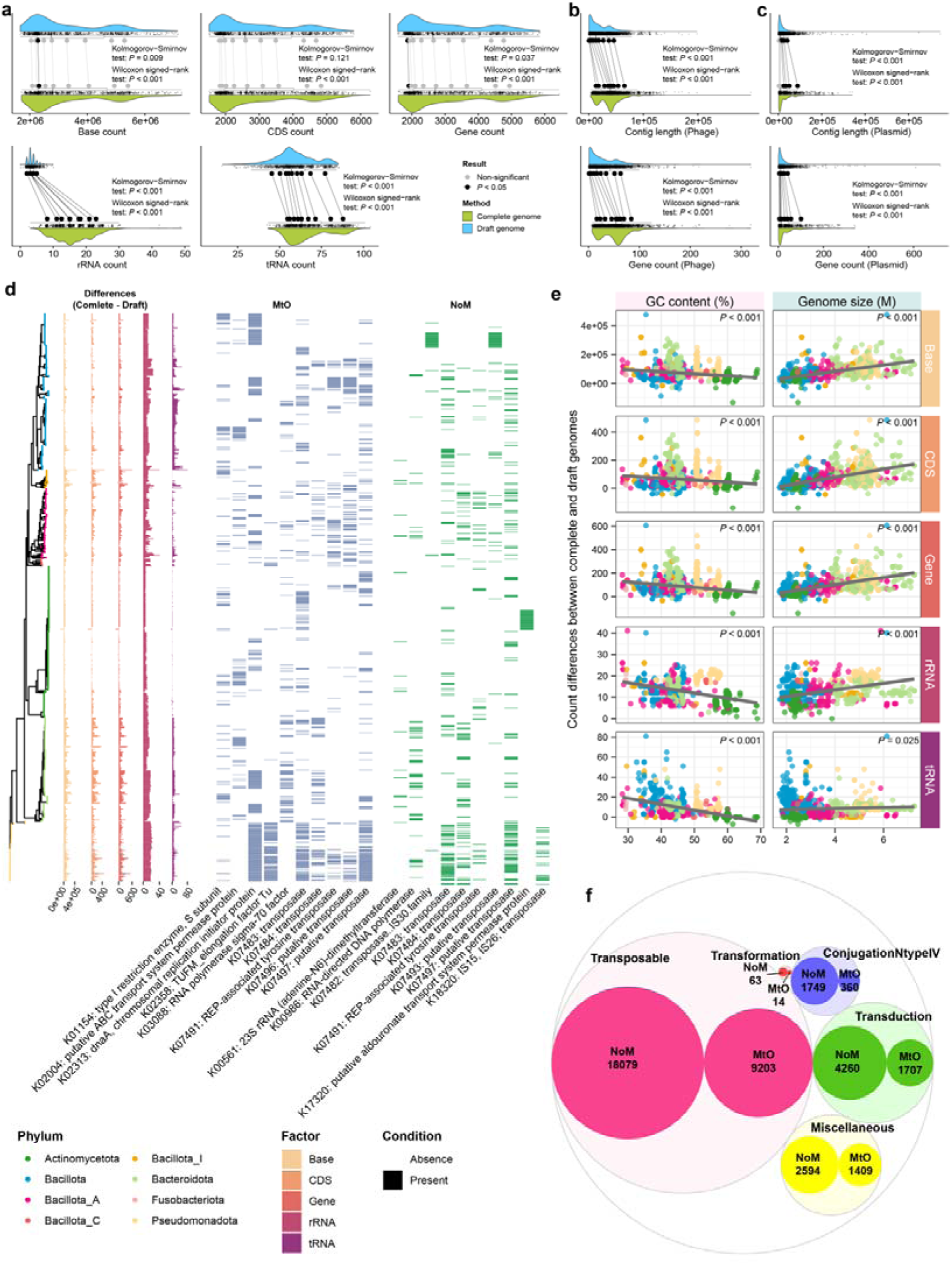
Complete genomes uncover a refined landscape of genetic elements. **a** The raincloud plots show the difference in the number of multiple genomic metrics, including base counts, CDS counts, gene counts, rRNA counts, and tRNA counts, between complete genomes and the corresponding draft genomes. **b** The raincloud plots show the difference in contig length and gene count of the phages between complete genomes and the corresponding draft genomes. **c** The raincloud plots show the difference in contig length and gene count of plasmids between complete genomes and the corresponding draft genomes. **d** Phylogenetic tree colored by bacterial phylum is shown on the left, barplots show the difference in the number of multiple genomic metrics between complete genomes and the corresponding draft genomes for each genomes, and the heatmap shows the KO annotation of MtO genes and NoM genes in each genome. **e** The linear relationships of the count difference of multiple genomic metrics between complete genomes and the corresponding draft genomes with GC content and genome size. **f** Circle packing plot displaying different HGT types associated with MtO genes and NoM genes, the color indicates HGT types, and the circle size indicates number of genes.

We also compared the detailed differences in CDSs (Supplementary Data4) showing that the complete genomes contained a large number of CDSs missing in the drafts (NoM) and CDSs with reduced copy numbers from the draft (NoM). Notably, we found that these genes associated with transformation, transduction, and conjugation and present in complete genomes were largely underrepresented in the draft genomes (Supplementary Fig. 6, Fig. 3d), indicating that complete genomes provided an accurate resolution of intricate HGT within bacterial genomes.

### Complete genomes resolve phage integration rules and auxiliary metabolic potential

Based on the chromosome-scale resolution of 1,150 complete genomes, we constructed a high-quality phage catalog (CCGRv) comprising 3,046 viral sequences (Supplementary Fig. 7a). This dataset represents a substantial leap in genomic integrity, as 60.09% of phages exhibited significant quality improvements compared to draft-derived assemblies, with the average completeness rising from 39% to 74% (Supplementary Fig. 7b-e, Fig. 4a). A striking example of this resolution is CCGRVR01473, a single complete phage sequence that successfully bridged eight fragmented contigs from the corresponding draft genome into a contiguous scaffold. This refined resolution enabled us to accurately delineate the diversity and integration patterns of gut phages. While phages from Pseudomonadota and Bacteroidota exhibited the highest richness, the phylum Bacillota harbored the majority of novel viral clusters (Supplementary Fig. 7f, Fig. 4b). Furthermore, we observed that while most phages were host-specific, a subset demonstrated potential cross-species transmissibility (Fig. 4c).

**Fig. 4.**
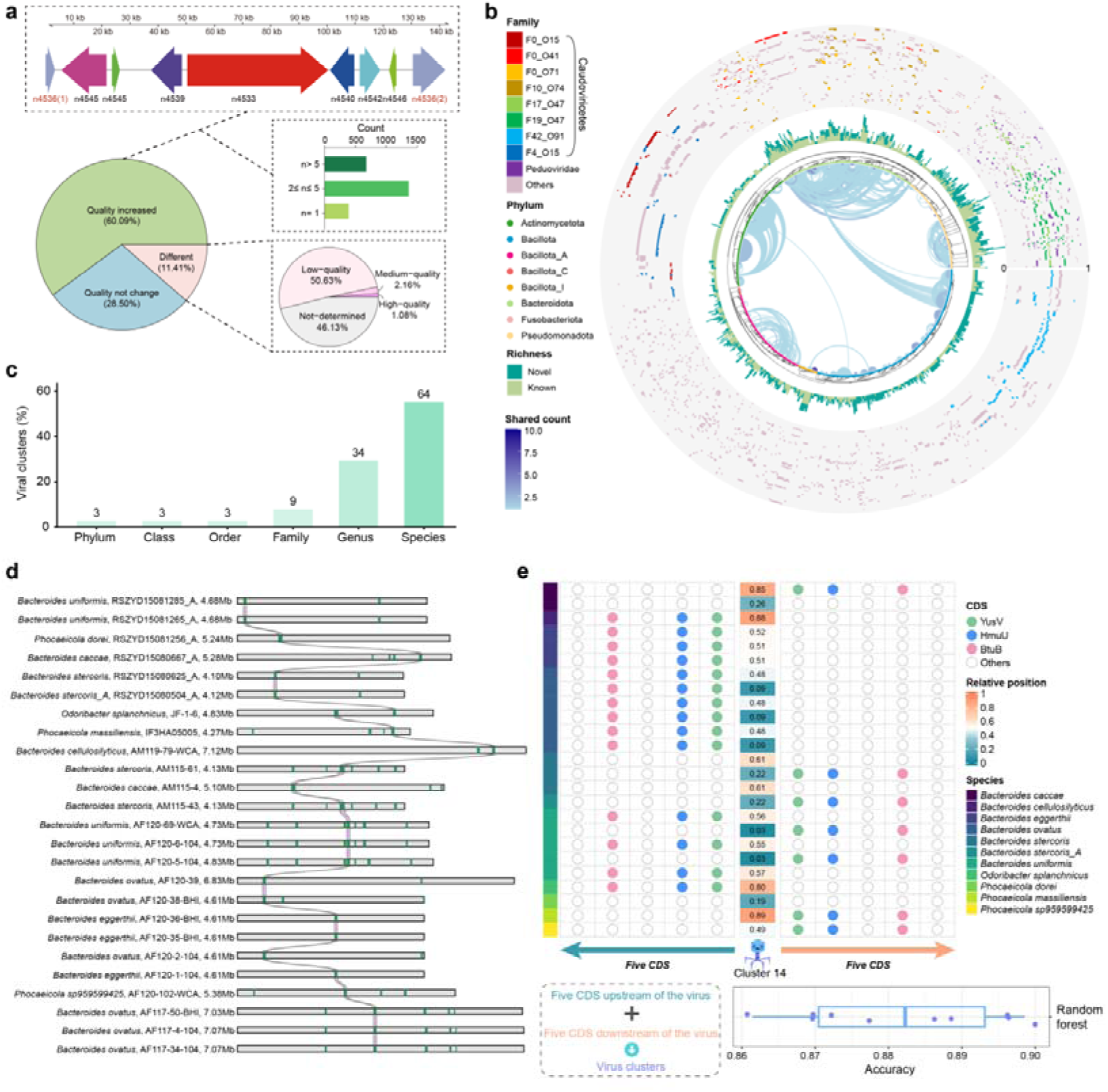
Complete genomes unveil the refined landscape of phage diversity and chromosomal integration hotspots. **a** Improvement in phage genome assembly quality. At the top is an example where sequences of CCGRVR01473 (the phage predicted from a complete genome) comprised 8 different phage sequences derived from a draft genome. The pie chart shows the length changes of phages from the draft and phages from the complete genomes. The barplot shows the counts of different numbers of phages from the draft genomes forming one phage in the complete genomes. **b** In the innermost circle, connections between host genomes represent the number of shared phages, indicating potential phage transfer events; in the second circle, node colors denote the taxonomic classification of host genomes at the phylum level; in the third circle, barplots illustrate the richness of phage clusters, including both novel and known phage clusters; and in the outermost circle, the heatmap shows the positions of phages on each genome. For ease of visualization and interpretation, relative positions were normalized to a 0-1 scale, where 0 corresponds to the location of *dnaA*. The bottom-left barplot displays the number of potential cross-phylum phage transfers, and the pie chart shows the proportions of cross-phylum versus within-phylum transfers among all potential transfer events. **c** Host specificity and cross-taxa conservation analysis. The bar chart displays the percentage of viral clusters conserved at different taxonomic ranks. **d** The chromosomal integration of phage cluster 14 across distinct host species. The green blocks/lines highlight the specific integration location of phages. **e T**he positions of phage cluster 14 on bacterial chromosomes and its upstream and downstream CDSs. The boxplot shows the prediction of phage clusters based on five CDSs upstream and downstream of each phage using a random forest model to demonstrate the sensitivity of phage positions. The whole procedure was repeated 10 times with independent seeds.

Of note, complete genomes allowed us to decipher the precise integration patterns of these cross-species phages. We focused on Phage Cluster 14, a broad-host-range group widely distributed across distinct species, such as *Bacteroides uniformis* and *Phocaeicola dorei*. Although the absolute genomic coordinates of these prophages varied significantly across different host chromosomes, our synteny analysis revealed that their integrations were governed by strict local genomic context (Fig. 4d). Specifically, the insertion sites of Phage Cluster 14 were consistently flanked by specific nutrient metabolism genes, such as the vitamin B12 transporter *Btu*B, *Hmu*U, and *Yus*V, regardless of the host species (Fig. 4e). The strict conservation of this flanking syntax was verified by a random forest model, which predicted all phage identity in our data solely based on these upstream and downstream CDS patterns with greater than 85% accuracy.

Beyond integration specificity, our analysis of auxiliary metabolic genes (AMGs) showed that phages actively contributed to host metabolic networks. We identified a broad repertoire of AMGs involved in cofactor, vitamin, and carbohydrate metabolism (Supplementary Fig. 8a, b). An example of phage-host synergy was observed in *Eubacterium ramulus*, where phages encoded key enzymes for nicotinate and nicotinamide metabolism (e.g., *nad*E, *pnc*A), effectively complementing the host’s pathway to enable robust NAD^+^ biosynthesis (Supplementary Fig. 8c). In summary, these findings emphasize that gut phages are not merely passive genetic passengers, but rather strategically integrated. They are anchored by specific functional genes and equipped with AMGs to synergistically regulate host adaptation.

### Plasmids drive strain-level metabolic heterogeneity and therapeutic efficacy

To explore the functional contribution of extrachromosomal elements to host fitness, we constructed a catalog of 1,124 plasmids (CCGRp) from our complete genomes. These plasmids exhibited extensive mobility, with inter-phylum transfer events primarily occurring between Bacteroidota and Bacillota (Supplementary Fig. 9a). These mobile elements are classically recognized as vectors for disseminating antimicrobial resistance, evidenced here by the multi-species clustering of aminoglycoside and tetracycline resistance genes (Supplementary Fig. 9b). We identified five plasmids, each harboring ≥5 antimicrobial resistance genes (ARGs) that were predominantly arranged into discrete gene clusters (Supplementary Fig. 9c). Moreover, we uncovered evidence for putative HGT of ARGs between bacterial chromosomes and plasmids. A representative example is a conserved ARG cluster encoding a drug metabolite transporter superfamily protein, a tetracycline repressor, and a tetracycline resistance gene—detected in both the chromosome of *Escherichia coli* and a plasmid of *Klebsiella pneumoniae* (Supplementary Fig. 9c, d). Our functional profiling revealed that these plasmids also served as expansive reservoirs for metabolic traits, particularly within the family *Lactobacillaceae* (Supplementary Fig. 10).

Prompted by this metabolic potential, we identified a specific plasmid in *Levilactobacillus brevis* carrying a conserved *scrK* gene which encodes an enzyme with fructose degrading ability (Fig. 5a). This plasmid was experimentally validated to confer a profound growth advantage when fructose was the sole carbon source, distinguishing Plasmid^+^ strains from their Plasmid^-^ counterparts in vitro (Fig. 5b-d). We next investigated whether this plasmid-mediated metabolic capacity could translate into therapeutic efficacy *in vivo*. In a high-fructose (HF, 10%) diet-aggravated dextran sulfate sodium (DSS) colitis model, colonization with Plasmid^+^ *L. brevis* significantly reduced fecal fructose concentrations (Fig. 5h), effectively scavenging the dietary overload. This metabolic intervention translated into a significant amelioration of the symptoms of DSS-induced colitis, evidenced by increased colon length, reduced disease activity, and suppressed systemic and colonic inflammation compared to Plasmid^-^ strain-treated mice (Fig. 5e-p). Furthermore, the Plasmid^+^ strain increased intestinal barrier integrity, preventing the HF-induced loss of MUC2 and tight junction proteins (Fig. 5q-u).

**Fig. 5.**
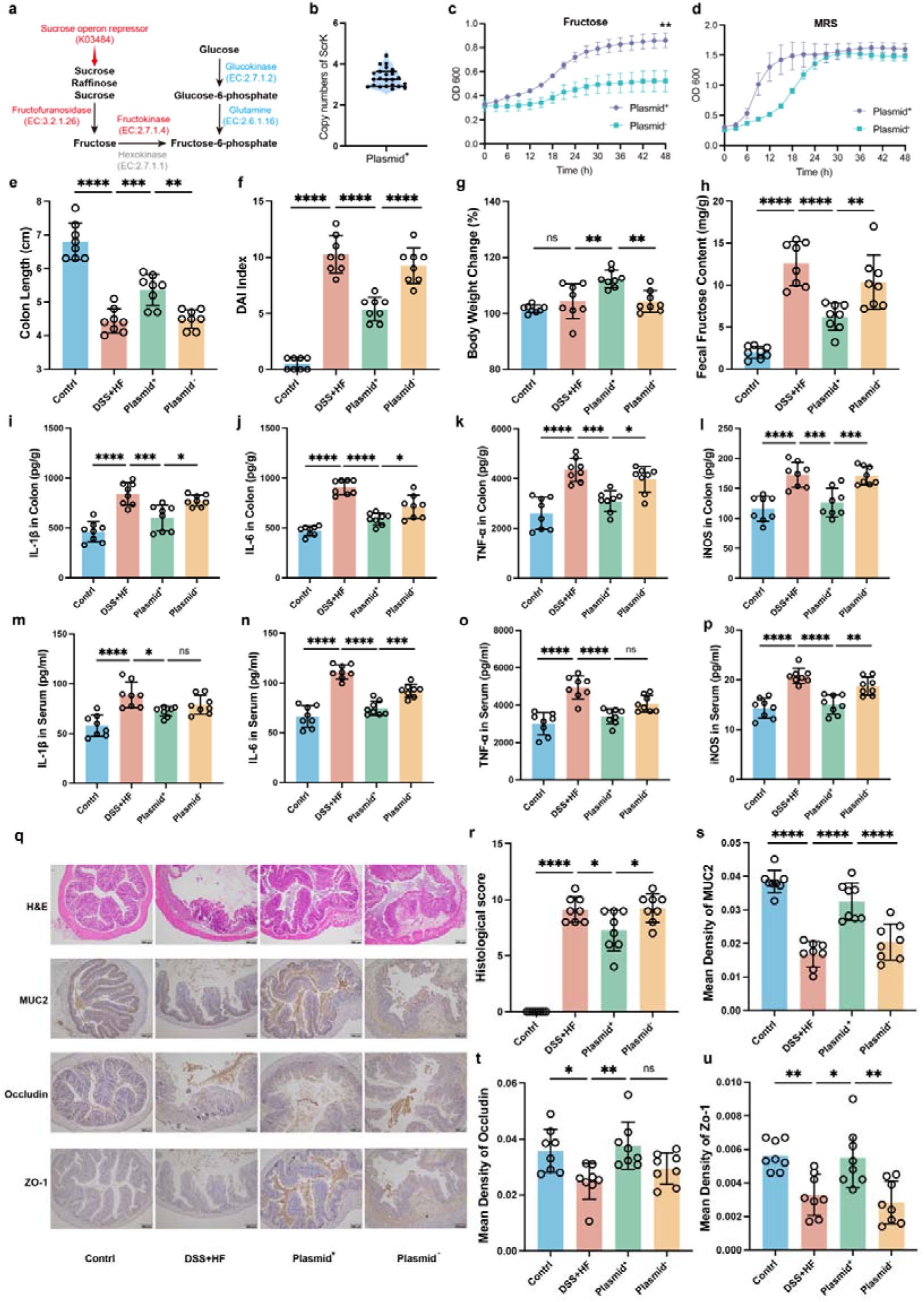
A plasmid-encoded fructose utilization gene empowers *Levilactobacillus brevis* to ameliorate high-fructose diet and DSS-induced colitis. **a** Schematic representation of the plasmid-borne metabolic pathway for fructose utilization. **b** The fold change of *scrK* gene copy numbers in two types of *L. brevis* strains (Plasmid^+^/Plasmid^-^). **c, d** Growth curves of Plasmid^+^ and Plasmid^-^ *L. brevis* strains cultured in minimal medium supplemented with fructose (c) or in MRS broth (d). **e-u** In vivo evaluation of the protective effects of *L. brevis* strains on DSS-induced colitis under a high-fructose (HF, 10%) diet. **e-g** Physiological assessments including colon length (e), Disease Activity Index (DAI) scores (f), and body weight change (g). **h** Fecal fructose content analysis indicating the fructose-scavenging capability of the Plsamid^+^ *L. brevis* strain in vivo. **i-p** Pro-inflammatory cytokine levels in colonic tissue (i–l) and serum (m–p), including IL-1β, IL-6, TNF-α, and iNOS. **q** Representative images of H&E staining (top row) and immunohistochemical staining for MUC2, Occludin, and ZO-1 in colonic sections. Scale bars, 200 µm. **r-u** Quantitative analysis of histological scores (r) and the mean optical density of MUC2 (s), Occludin (t), and ZO-1 (u). Data are representative of three independent experiments and presented as mean ± s.e.m. in c–u. Statistical significance was determined using two-way ANOVA (c, d) or one-way ANOVA followed by Tukey’s post hoc test (e–u): \**P* < 0.05, \*\**P* < 0.01, \*\*\**P* < 0.001, \*\*\*\**P* < 0.0001; ns, not significant.

To confirm that the observed effects were causally linked to fructose scavenging rather than general probiotic effects, we repeated the experiment under a low-fructose (LF, 1%) diet. In the absence of fructose excess, the therapeutic advantage of the Plasmid^+^ strain was completely abolished, with no significant differences observed in colitis symptoms, cytokine levels, or barrier markers compared to the Plasmid^-^ or vehicle groups (Supplementary Fig. 11a-q). Collectively, these data demonstrate that complete genomes enable the resolution of plasmid-encoded metabolic modules, such as the *scrK* gene that drives specific, strain-level therapeutic outcomes.

### Mobile genetic elements drive the expansion and strategic trade-offs of host defense systems

Beyond their metabolic contributions, MGEs exert profound selective pressure on host genomes, driving an adaptive “arms race.” Leveraging our complete genomes, we systematically mapped the landscape of defense and anti-defense systems (Supplementary Fig. 12a-c). We observed a significant positive correlation between the abundance of defense systems and the density of MGEs (phages and plasmids) within a genome (Supplementary Fig. 12d), confirming that MGE burden is a driver of host immune expansion. Interestingly, this co-evolutionary dynamic exhibits distinct strategies depending on genomic plasticity. In “open” genomes with a low core gene fraction, defense systems expand rapidly to counter MGE influx; conversely, in evolutionarily “conserved” genomes (core gene fraction > 50%), we observed a negative correlation between defense system abundance and genomic conservation (Supplementary Fig. 13e, Supplementary Fig. 14). This suggests a fundamental trade-off where bacteria must balance the fitness costs of maintaining extensive immune arsenals against the benefits of genomic stability.

### Distribution of plasmids and phages relative to human health and disease

We calculated the distributions of phages and plasmids in healthy cohorts from China, and the Netherlands, and in the HMP (Fig. 6a). Phages with members of the phyla Bacillota_A and Bacteroidota as hosts exhibited high prevalence and abundance, whereas this was only observed for a subset of plasmids hosted by Bacteroidota. As expected, there were significant differences in the composition of phages and plasmids among the three cohorts, reflecting the potential influence of geographic origin (Supplementary Fig. 14). Some phages and plasmids were unique to Chinese (Supplementary Fig. 15). We also found that phages and plasmids with high prevalence predominantly hosted by genomes from the phylum Bacteroidota and Bacillota_A exhibited distinct differences relative to the occurrences of diseases (Fig. 6b). Thus, specific MGEs displayed strong correlations to liver cirrhosis (LC), atrial fibrillation (AF), and ankylosing spondylitis (AS). Conversely, in general, MGEs were showed less correlation to coronavirus disease 2019 (COVID-19) and essential hypertension (EH), and few correlations to inflammatory bowel disease (IBD) and colorectal cancer (CRC). Intriguingly, the abundances of a large proportion of phages and plasmids were markedly increased in AF group but decreased in AS group (Fig. 6b). Our results suggested that most of the phages and plasmids such as Phage Cluster 1023 positively correlated with AF encodes the toxin-antitoxin systems, such as MazF-MazE, which may promote the development of the disease by enhancing the virulence of bacteria. In addition, Plasmid Cluster 298 that was strongly negatively correlated with AS harbours the resistance-related gene ermC. We also found differences in the abundances of plasmids and phages between urban and rural areas (e.g. Phage Cluster 1273, Phage Cluster 1297, Plasmid Cluster 132, and Plasmid Cluster 298), suggesting an impact of the living environment. Notably, the majority of these biomarkers were novel, suggesting that our study enabled the discovery of microbial dark matter with potential biological impact and value. We observed overwhelmingly synergistic relationships between phages and plasmids, with only a minor proportion of negative correlations (Fig. 6b). We next investigated whether the phages and plasmids could show co-evolution patterns with their hosts (***See details in Methods***). The results showed that in 88.89% of the selected species, the phages and plasmids tended to significantly co-evolve with their hosts (Supplementary Fig. 16). For example, *Parabacteroides distasonis* genomes clustered into four distinct lineages in the phylogenetic tree, each harboring fixed combinations of plasmids and phages (Fig. 6c). This suggested that these elements might facilitate adaptive evolution within the host. Moreover, we developed a random forest classifier to distinguish disease cases from healthy controls using the biomarkers identified from each disease cohort. These models demonstrated moderate to excellent diagnostic performance in AF, AS, schizophrenia (SCZ), and LC, with median area under the receiver operating curve (AUROC) > 0.7 (Fig. 6d). These findings indicate that the selected phages and plasmids with specific functional genes could represent new microbiome signatures associated with diverse human diseases.

**Fig. 6.**
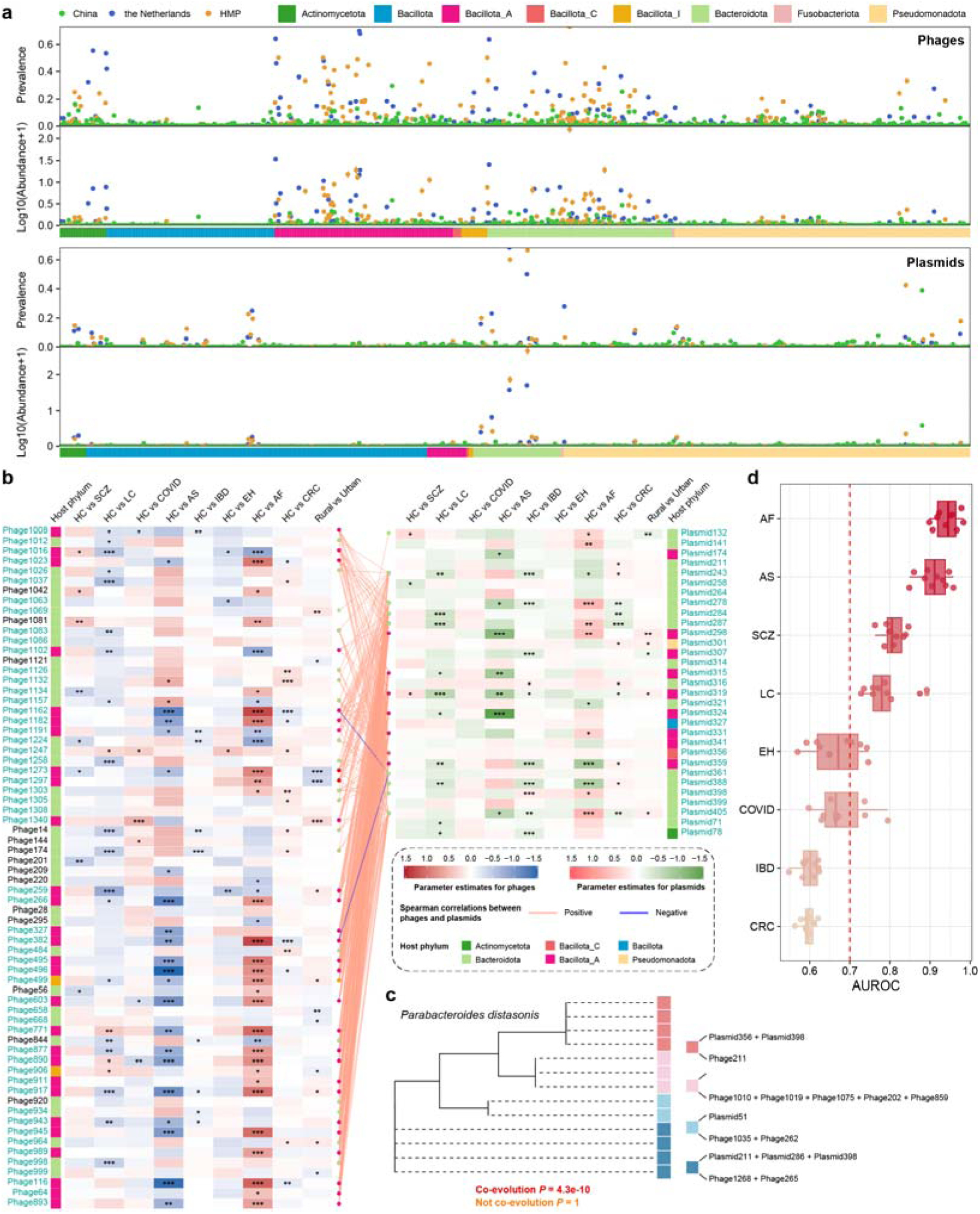
Distributions of high-quality plasmids and phages from completed genomes and their associations with disease development. **a** Abundance and prevalence of plasmids and phages in healthy cohorts of China, HMP, and the Netherlands. **b** Differences in the abundances of plasmids and phages between healthy and disease states shown by slopes of regression fits based on linear regression models. c The co-evolution of the genomes in Paraba*cteroides distasonis* with their phages and plasmids. **d** The performance of random forest classifiers to distinguish disease cases from healthy controls tested by AUROC. Asterisks indicate statistical significance (*** *P* < 0.001, ** *P* < 0.01, and * *P* < 0.05).

## Discussion

The draft genomes impede our in-depth exploration of the genomic information and biological characteristics of gut microbiota, but complete genome collection for gut bacteria is still lacking. In this study, we have obtained 1,150 circularized bacterial genomes, which made a great contribution to current public databases. The complete genomes provide information for us to investigate the gene placement patterns of bacterial orthologs. Our results showed that the complete genomes significantly increased the number of 16S rRNA genes and tRNA genes and performed much better at plasmid and phage prediction compared to drafts. Moreover, 53% of plasmid sequences derived from the draft genome mapped to the chromosome in the complete genomes, suggesting that plasmid prediction tools may overestimate the number of plasmids in previously fragmented draft genomes. Notably, we found many genes, especially for MGE associated genes, were missing or only presented as a single copy in the draft genomes, implying the CCGR reveal more refined mobile element-host interactions.

The location of functional genes and phages on chromosomes is crucial for understanding gene expression and further explaining the infection mechanism of phages. Previous studies have demonstrated that roughly two-thirds of bacterial genes exhibit positional preferences, with genes closer to the replication origin having higher intracellular copy numbers during rapid bacterial growth^32,34^. Our results demonstrate that bacterial genomic organization is not governed by a monolithic biophysical law, but by divergent evolutionary strategies. Bacillota relies on rigid topological constraints, strictly sequestering long genes to the leading strand. However, this strategy proves brittle; rapid growth correlates with increased genomic fragmentation, suggesting that strict conflict-avoidance mechanisms become overwhelmed by the mechanics of fast replication. In contrast, Pseudomonadota exhibits architectural robustness. By decoupling gene length from strand specificity and maintaining structural order even during rapid growth, these lineages appear to possess superior conflict-resolution machinery that allows them to manage rather than strictly avoid collisions.

Our study reveals that phages and plasmids are not merely genomic passengers but potent drivers of disease pathophysiology, acting as robust biomarkers through specific functional contributions. Beyond well-known virulence factors like toxin-antitoxin systems and antibiotic resistance genes, our complete genome-resolved analysis uncovers a deeper layer of metabolic synergy.

Crucially, we demonstrate that MGEs are primary architects of strain-level metabolic heterogeneity. While excessive fructose intake is a known driver of colitis and metabolic dysregulation, we identified a plasmid-encoded *scrK* gene in specific *L. brevis* strains that confers a scavenging phenotype. This plasmid-dependent adaptation enables the host to rapidly metabolize dietary fructose, thereby mitigating fructose-induced intestinal barrier damage and inflammation^35–41^. This finding exemplifies a critical paradigm: functional differences between strains could be dictate by the dynamic accessory genome rather than the core chromosome. Consequently, probiotic efficacy and pathogenicity may be misattributed in species-level analyses that overlook these mobile traits.

Furthermore, the strict chromosomal integration hotspots of phages and their carriage of AMGs reflect a long-term co-evolutionary strategy to modulate host fitness. These synergistic interactions enhance the reliability of MGEs as disease markers, as their presence signifies specific functional demands of the gut environment. Collectively, our work underscores that moving from fragmented drafts to complete genomes is indispensable. Only by resolving the precise linkage between MGEs and their hosts can we decode the hidden functional diversity that defines the personalized nature of the human gut microbiome and its impact on health.

## Materials and Methods

### Sample collection and cultivation of bacterial isolates

Fecal samples were collected from healthy volunteers not taking any antibiotics in the last six months prior to sampling or suffering from intestinal diseases. The sample collection was approved by the Institutional Review Board of BGI Ethical Clearance under number BGI-IRB 22112-T1. Fecal samples were diluted and spread on agar culture mediums under anaerobic conditions. We then picked thousands of single colonies and transferred each of them to 2 ml of liquid medium for further culture. We used 16S rDNA PCR to preliminarily identify the species, a diverse range of species were selected for subsequent genome sequencing.

### Genome sequencing and genome assembly

A hybrid assembly approach integrating DNBSEQ short-reads and CycloneSEQ long-reads was used to obtain bacterial complete genomes. The method of whole-genome sequencing was as described by Liang et al^42^. The DNA of each strain was extracted using the Magen MagPure DNA Kit for high-throughput applications, the CycloneSEQ library preparation and sequencing followed the manufacturer’s guidelines and protocols. The libraries were quantified using a Qubit fluorometer and sequenced on the CycloneSEQ WT02 platform according to sequencing protocols. The method of genome assembly was as described by Liang et al^42^. Long-read data was filtered using NanoFilt^43^ with parameters “-q 10 -l 1000” to retain reads longer than 1,000 bp and with a quality score greater than Q10. Short-read data was processed using Fastp^44,45^ with default parameters, except the length requirement was set to 50. Short-read assembly was performed using Unicycler with only the short reads ‘-1’ and ‘-2’ as input, and the ‘-depth_filter’ set to 0.01 to remove low-depth contigs^31^. For hybrid assembly, the same ‘depth_filter’ of 0.01 was used, with the addition of ‘-l’ long reads as input, while all other parameters were set to default. For the assembly of MAGs from 127 contaminated culture isolates, flye v2.9.1-b1784^46^ with the ‘--meta’ options was used to assemble the preprocessed clean long reads, then short-read polishing was performed using Nextpolish v1.4.1^47^ with default parameters.

### Genome quality assessment and gene prediction

The circularity of each genome was evaluated using the assembly results from Unicycler and Flye^31,48^. Genome quality was assessed using CheckM2, circular genomes with >95% completeness and <5% contamination were retained for further analysis^49^. To perform a standard processing of circular genomes for subsequent gene placement analysis, the “fixstart” function of Circlator (v1.5.5) was used to rotate the largest circular fragment (here we refer to as the main circular chromosome) of each circular genome, to ensure the *dnaA* gene positioned as its origin^50^. This process was implemented for all genomes except for four genomes in which the *dnaA* gene was absent or not located on the main circular chromosome. Moreover, for the main circular chromosome with multi-copy *dnaA* genes, we used the information of GC skew to manually select a correct *dnaA* gene as the origin.

### Collection and process of genomes from public dataset

To assess the contribution of the CCGR database to the currently available human gut bacterial complete genomes, bacterial complete genomes were downloaded from the NCBI RefSeq database (09-2024) using the ncbi-genome-download tool. Genomes assembled at complete level were selected, the atypical genomes, genomes from large multi-isolates, and metagenome-assembled genomes (MAGs) were excluded. Subsequently, a total of 1,562 human gut bacterial genomes passing CheckM2 quality control with the threshold of >95% completeness and <5% contamination were retained. The 1,562 genomes and 1,150 genomes of the CCGR dataset were clustered at species level using dRep (v3.5.0) with a 95% average nucleotide identity (ANI) threshold (-sa 0.95 -nc 0.30) to identify common and unique species^51^.

### Phylogenetic and taxonomic analyses

All genomes were taxonomically annotated using GTDB-Tk (v2.4.0) ‘classify_wf’ function against the GTDB database r220^52^. The genome maximum-likelihood phylogenetic tree was constructed using ‘infer’ function of GTDB-Tk based on 120 bacterial marker genes and visualized by iTOL^53^.

### Gene placement analysis

To enable global positional comparisons of genes across genomes of varying sizes, we normalized genomic coordinates relative to the replication origin (defined by *dnaA*) and terminus (defaulted to the genomic midpoint).

First, genomic data were collapsed to the species level by calculating the median normalized position for each orthologous gene to mitigate sampling bias. A “Consensus gene rank” was established for each phylum by ordering genes (present in > 50% of species) based on their median positions. Global synteny was visualized by plotting this consensus rank against the actual normalized position. Second, we quantified three key metrics to characterize genomic architecture: (1) Strand switch rate, defined as the frequency of transcriptional orientation changes between adjacent genes, calculated as the percentage of gene pairs with opposing orientations relative to the total gene count, serving as a proxy for genomic fragmentation and the extent of co-directional gene organization; (2) Leading strand bias, calculated as the percentage of genes encoded on the leading strand based on replichore geometry (forward strand for the right replichore, reverse strand for the left); and (3) gene-level distributions of switch rate and strand, where we analyzed the frequency distribution of them for individual genes to characterize phylum-specific strategies. It should be noted that, in order to ensure the reliability of the results, we only conducted these analysis on the five phyla with the largest number of genomes here.

To evaluate the constraints imposed by physical gene length and function, genes were binned by coding sequence length (log-transformed), and the lagging-to-leading length ratio was calculated for each genome. Functional constraints were assessed by mapping proteins to Clusters of Orthologous Groups (COGs), with architectural metrics calculated separately for each category.

Finally, to link genomic architecture with organismal growth dynamics, we predicted maximal growth rates from genomic sequences using gRodon^54^. We correlated these predicted growth rates with genomic architectural metrics. Specifically, we analyzed the relationship between growth rate and the global strand switch rate, the leading strand bias of genes within the dnaA-proximal region (1/3 of the replichore), and the accumulation of genes in high-risk zones, defined as regions proximal to *dnaA* but encoded on the lagging strand where replication-transcription conflicts are most severe.

### Evaluating the improvement of complete genomes in comparison with draft genomes

To assess the enhancement of complete genomes over draft assemblies in terms of different genomic features, the number of bases, 16S rRNA genes, tRNA genes, protein-coding sequences, and total genes were gathered from the prediction results of Prokka (v1.14.6) for both 966 complete genomes and their corresponding draft genomes^55^. For further comparison of CDS, we used MMseqs2 (15.6f452) linclust function (--cov-mode 5 -c 1 --min-seq-id 1 --kmer-per-seq 80) to identify matched and unmatched CDS between complete genomes and draft genomes^56^.

### Reconstruction of plasmid and phage catalog

In general, the genomes of plasmids and phages were predicted using geNomad (v1.11.0) with the parameters (end-to-end, --cleanup, --default)^57^. CheckV was used to assess the quality of phage genomes^58^. For clustering of plasmid and phage sequences, CheckV built-in script was used 95% ANI and 85% coverage as criteria, and sequences larger than this criterion were classified into a unified cluster. We determined that plasmids and phages from the same cluster could potentially invade the same host.

For the novel plasmids and phages, we respectively used representative plasmid collections from PLSDB (2024_05_31 version) and representative phage collections from multiple databases provided by UHGV as the references, and used a mash distance of 0.05 as the cutoff for whether a plasmid/phage was novel^59–61^.

To identify plasmids that provide unique functions relative to the bacterial chromosome, we used Diamond to align the predicted CDSs of each plasmid^62^.

### Blastn was used to find the quality increase of phages and plasmids

For quality improvement, we used a longer than 5% increase in genome length as the criterion. There are a total of 4830 plasmid sequences from the short-read genome assembly, of which 3278 sequences are aligned to the main loop sequence of the genome completion map (E-value < 10e-5). Among these, 2571 short-read genomic-derived plasmid sequences had ≥95% alignment coverage to the chromosomes. We defined these plasmids as misidentified. For plasmids that failed to align with the main circle sequence, we aligned them with plasmids derived from the complete genomes. For phages, we found that 60.09% of the sequences had a length increase of more than 5%. 28.5% of the sequences were identified as unchanged and 11.41% of the sequences were identified as no mapping.

### Prediction of defense and anti-defense genes/systems

The anti-MGE defense genes/systems of bacterial genomes predictions were performed using defensefinder (V2.0.0) based on the database (defense-finder-models: 2.0.2)^63^. We also investigated defense and anti-defense systems encoded on phages and plasmids. Briefly, phages and plasmid sequences predicted by geNomad were used to perform defensefinder predictions identical to those described above.

### Phage-encoded AMGs annotation and definition

Phage-encoded AMGs were annotated using VIBRANT (v1.2.1, default)^64^. According to the previous definition^65^, all obtained AMGs were divided into two categories: Class I AMGs refer to genes involved in the “metabolic pathways” defined in the KEGG database, while genes belonging to other pathways are considered class II AMGs.

### Comparison of the distribution of representative clusters of phages and plasmid in healthy and disease individuals in cohort

We used three natural population cohorts (4D-SZ, Netherlands cohort, HMP cohort) to explore the global distribution of our representative plasmid and viral sequences and used nine disease cohorts to explore the relationship between phages and plasmids and health (Detailed cohort information can be found in “Data Availability”). Specifically, we used representative phage and plasmid sequences to construct genome indexes for read mapping and aligned them by bowtie2^48^. In the calculation of the occurrence, in order to avoid false positives, we used a higher threshold to define whether the sequence appears in the sample as described in previous studies^66,67^. For plasmids, we used CoverM^68^ to define more than 50% coverage as occurrence, and for phages, we defined more than 75% coverage as occurrence (--min-read-aligned-percent 90 --min-read-percent-identity 90 --contig-end-exclusion 75 -m covered_fraction). To balance the effects of different lengths of phage or plasmid genomes on abundance, we used a TPM-like method to quantify abundance.

### The fructose degradation by *Lactobacillus brevis* strains

In order to explore the ability of different types of *L. brevis* strains to degrade fructose in vitro, we designed the following experiment. We have 26 strains and genomes of *L. brevis*, of which 23 are predicted to carry the plasmid (CCGRPL00051, Cluster 180) and 3 are predicted to not carry this plasmid. Due to time constraints, we were unable to successfully resuscitate all strains, so we randomly selected three resuscitated strains with plasmids and three strains without plasmids for the experiment. We designed a culture medium with fructose as the sole carbon source to verify the degradation status of different strains growing under this condition (Supplementary Data 8). The strains were activated on standard MRS medium and then cultured in liquid medium under different culture conditions. Then, we measured the OD value every 6 h and plotted the growth curve of the strain.

### Animal experiments and experimental design

Specific pathogen-free (SPF) male C57BL/6 mice (6-8 weeks old) were housed under standard conditions. To evaluate the protective effects of *Levilactobacillus brevis* strains driven by the fructose-metabolizing plasmid, mice were randomly divided into groups (n=8 per group). For the high-fructose (HF) model, mice were administered 10% (w/v) fructose in food. For the low-fructose (LF) control, mice received 1% (w/v) fructose. In both models, mice were orally gavaged daily with 10^9^ CFU of either Plasmid^+^ or Plasmid^-^ *L. brevis* resuspended in PBS. The vehicle group received PBS only. Following a 7-day bacterial colonization period, colitis was induced by adding 1.5% (w/v) dextran sulfate sodium (DSS, MW 36-50 kDa) to the drinking water for 7 days, followed by a 3-day recovery period with normal water.

### Assessment of colitis and sample collection

The Disease Activity Index (DAI) was monitored daily, assessing body weight loss, stool consistency, and rectal bleeding. At the endpoint, mice were sacrificed, and colon lengths were measured. Distal colon tissues were fixed in 4% paraformaldehyde for histological analysis (H&E staining) and immunohistochemistry (IHC) targeting MUC2, Occludin, and ZO-1.Fecal samples were collected to quantify fructose content using a Fructose Assay Kit according to the manufacturer’s instructions. Pro-inflammatory cytokines (IL-1β, IL-6, TNF-α, and iNOS) in colonic homogenates and serum were quantified using ELISA kits.

### Random forest models

We first employed a random forest model to predict phage clusters with five upstream and downstream CDSs around the phages serving as features. The model was also used to classify case and control from 8 kinds of disease cohorts, with the biomarkers serving as features. All the analysis was conducted using the R package randomForest and caret. Model training and hyperparameter tuning was performed on 70% of the data using a 5-fold cross-validation, while the other 30% was used for testing with the best hyperparameter setting. The whole procedure was then repeated 10 times with independent seeds. Model performance was evaluated using the AUROC.

## Supporting information

SFigures

STables

## Data availability

Our complete bacterial genomes and high quality plasmid and phage genomes have been uploaded at CNSA (CNP0007680). The numbers of the public cohorts we used in this study are as follows: Ankylosing spondylitis (ERP111669), Schizophrenia (ERP111403), hypertension (ERP023883), Cirrhosis (ERP005860), COVID-19 (DRP009646), Atrial fibrillation (ERP110580), Rural and urban (SRP237559), CRC (ERP012177, DRP004793, SRP320766), Chinese cohort (a part of 4D-SZ) (https://db.cngb.org/search/project/CNP0000426/), HMP (https://portal.hmpdacc.org/), the Netherlands cohort (https://ega-archive.org/studies/EGAS00001005027).

## Acknowledgements

This study is supported by National Key Research and Development Program of China (2025YFA1310200), Fundamental and Interdisciplinary Disciplines Breakthrough Plan of the Ministry of Education of China (JYB2025XDXM608), National Natural Science Foundation of China (82130068), Shenzhen Science and Technology Program (No. KCXFZ20240903094006009, No. JCYJ20241202124801003, SYSPG20241211173844007, SYSPG20241211173845014). We also thank the colleagues at BGI-Shenzhen for sample collection, DNA extraction, library construction, and sequencing.

## Author contributions

Y.Zou., L.X., H.Z., F.Z. and Y.D. supervised and led the project. Y.Z. initiated the idea and conceived the study. H.W., Y.G. and X.Y. designed and executed the analysis of phages, plasmids, defense systems and cohort study. W.H. and Y.G. designed and executed the analysis of genome localization and function. H.L., M.W. and Z.W. made functional annotation and download the cohort data. J.Y., Y.Zhong., J.M. and H.Z. preparation and cultivation of strains for sequencing. L.Y., Y.X. and J.Z. download the previous plasmid and phage database. T.Z., Y.Zhang. constructed the methods of assembly. Y.W., J.W., X.R., Y.F., X. T., Chuan Liu., X.S., B.W., X.J., Chuanyu.Liu., X.X., J.X., S.L., N.H. and P.G. collected information and interpreted data. J.Y., F.L., Z.Y., and W.L. constructed bacterial sequencing library. Y.Zou., L.X., H.Z. and K.K. reviewed and revised the manuscript. H.W., Y.G. and W.H. wrote the manuscript. All authors reviewed and approved the final version of the paper.

