## Supplementary material for "Complete Genomes of Cultivated Gut Bacteria Reveal Mobile Genetic Element-Driven Functional Diversity with Therapeutic Implications": SFigures

**Supplementary figures**

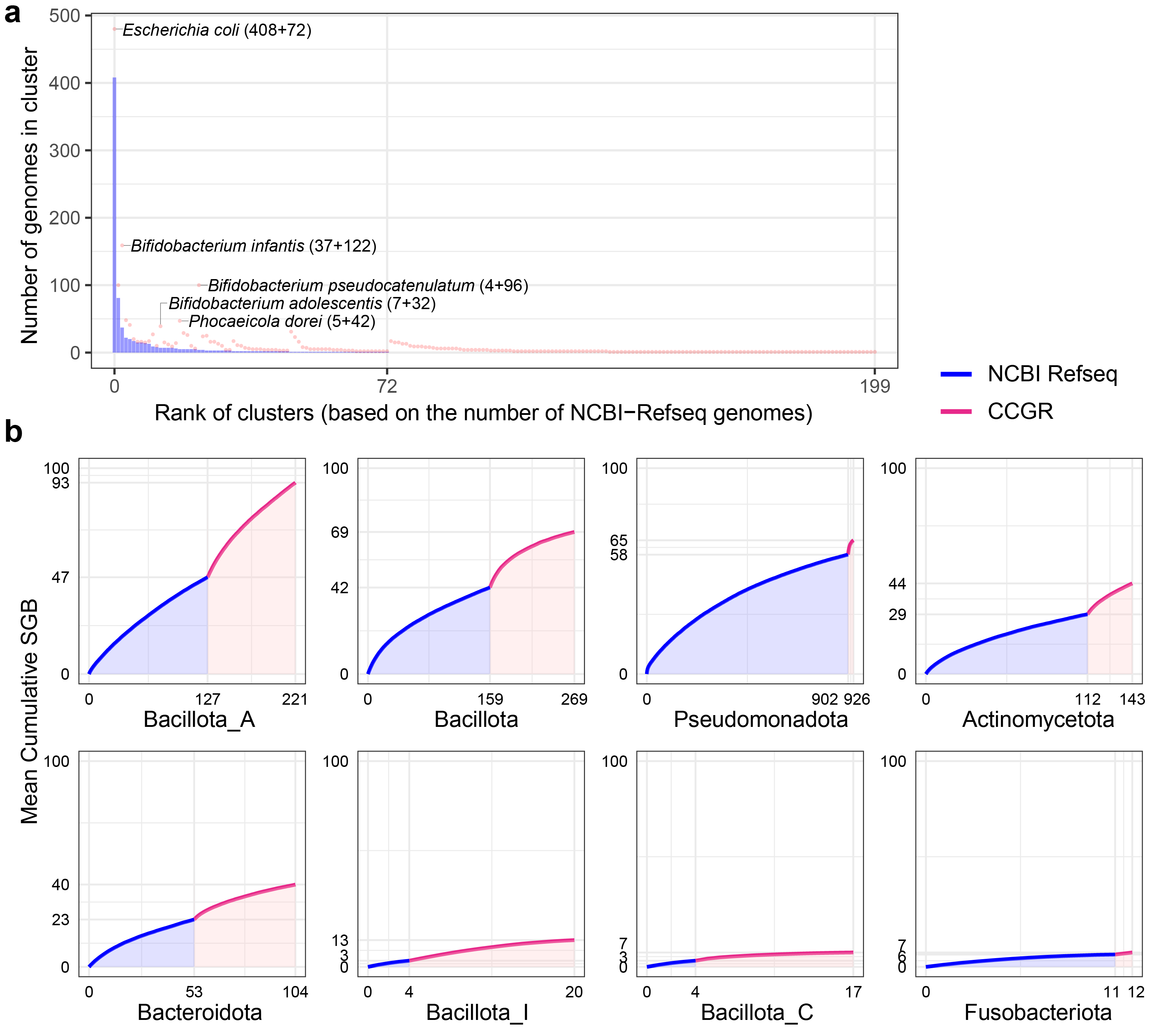

**Fig S1. Contribution of CCGR to the NCBI-Refseq database. a** Rarefaction curves of the number of representative clusters obtained as a function of the number of genomes analyzed, showing the contribution of CCGR to the NCBI-Refseq database across different bacterial phyla. **b** Contribution of CCGR genomes to the human gut bacterial complete genomes from NCBI-Refseq database. Clusters were sorted based on the number of the NCBI-Refseq genomes. Blue bars represent the distribution of NCBI-Refseq genomes in each cluster, and red dots represent the distribution of CCGR genomes in each cluster.

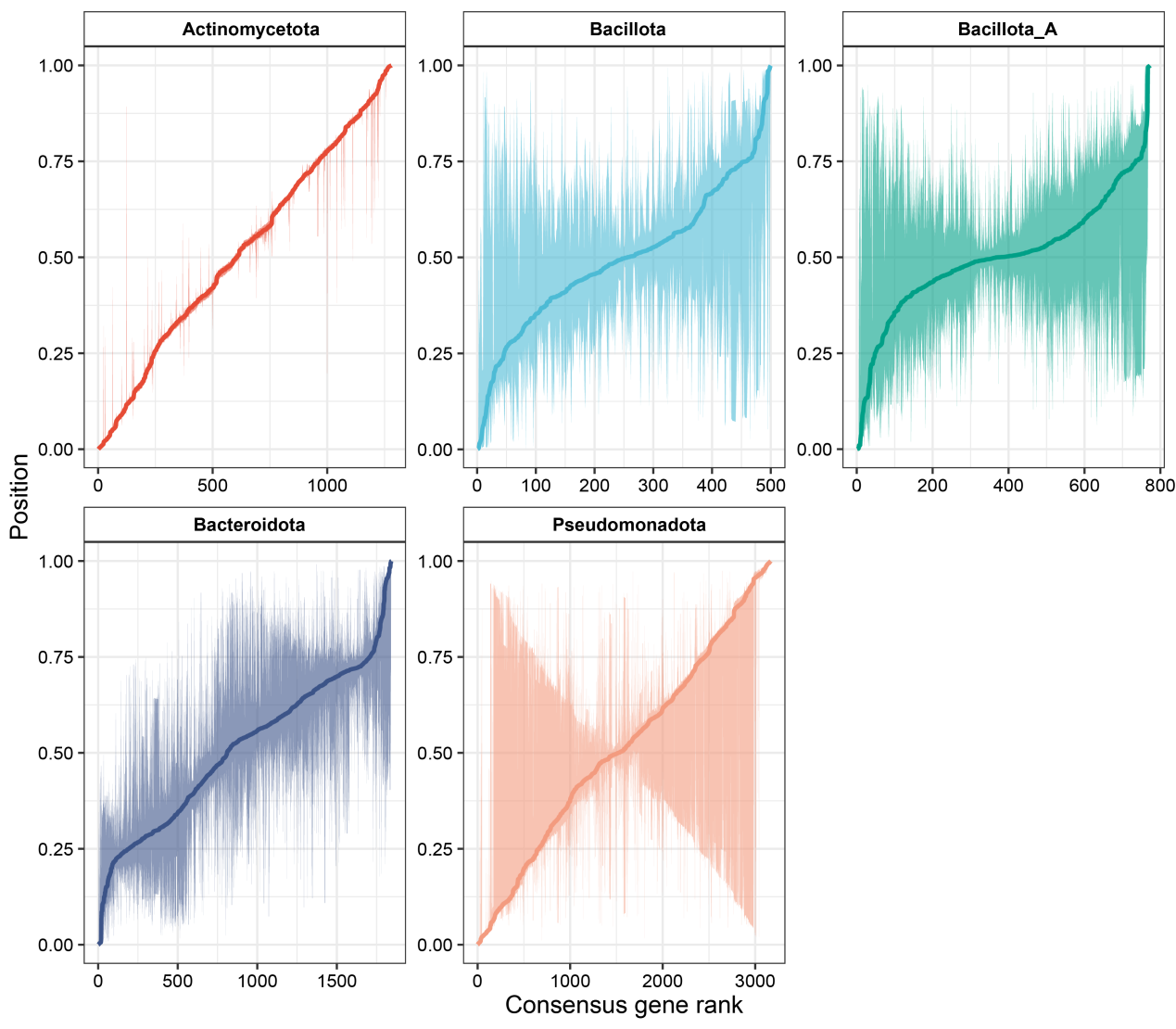

**Fig S2. Phylogenetic conservation of genomic architecture across phyla.** The X-axis represents the idealized order of genes with occurrence rate > 50% derived from the median position of each gene across all species within the major bacterial phylum. The Y-axis represents the actual normalized genomic position of each gene in individual genomes. For each gene, the normalized genomic position was calculated as the relative distance from *dnaA* (ranging from 0 to 1, where 0 and 1 represent dnaA and 0.5 represents the terminus). The solid line traces the median position across all genomes, while the shaded ribbon represents the interquartile range, indicating the extent of position variation or rearrangement. For each gene, the normalized genomic position was calculated as the relative distance from dnaA (ranging from 0 to 1, where 0 and 1 represent dnaA and 0.5 represents the terminus).

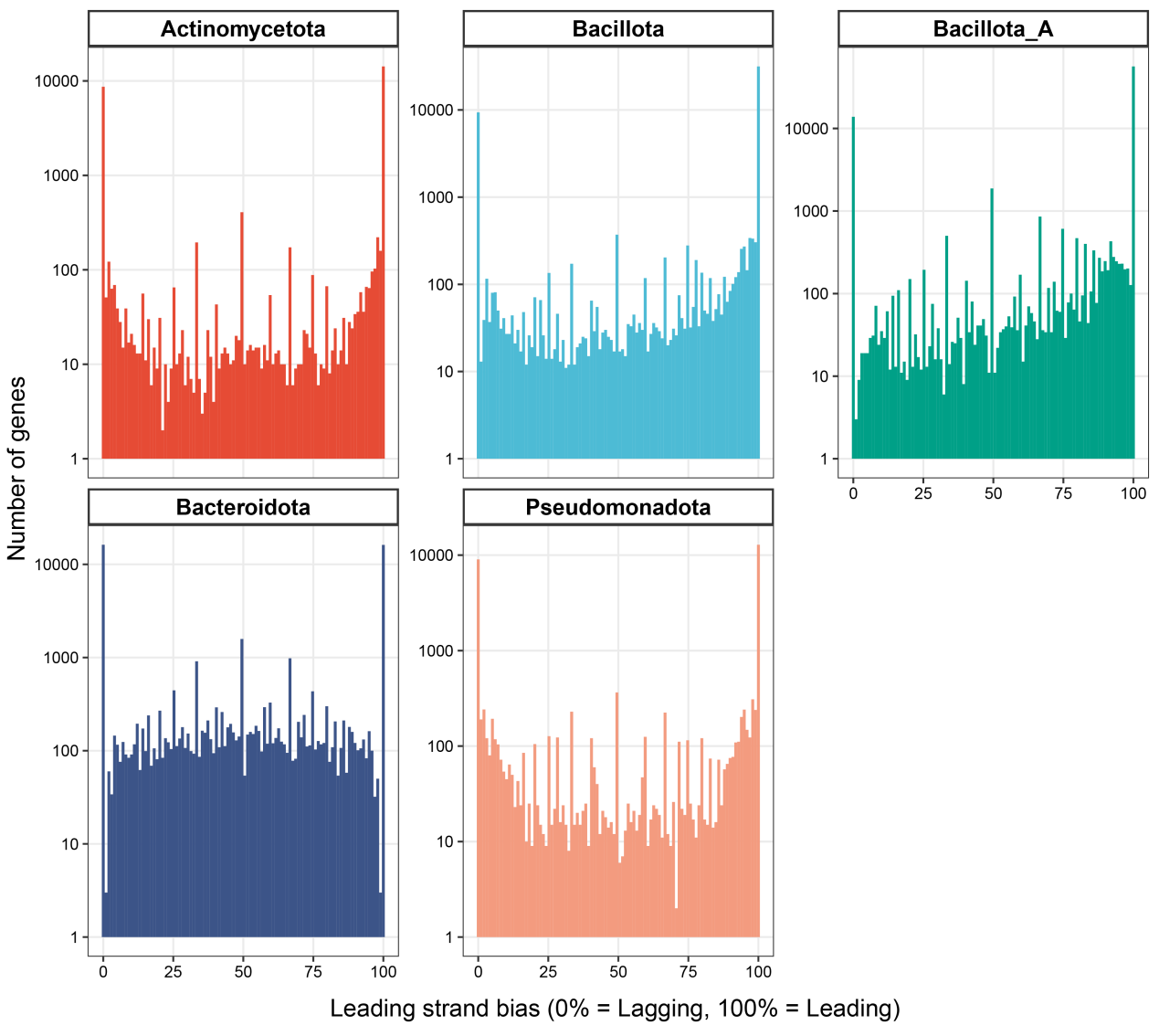

**Fig S3. Distribution of the leading strand bias for each gene across phyla.** The density histograms showing the leading strand bias of the genes within the major bacterial phylum.

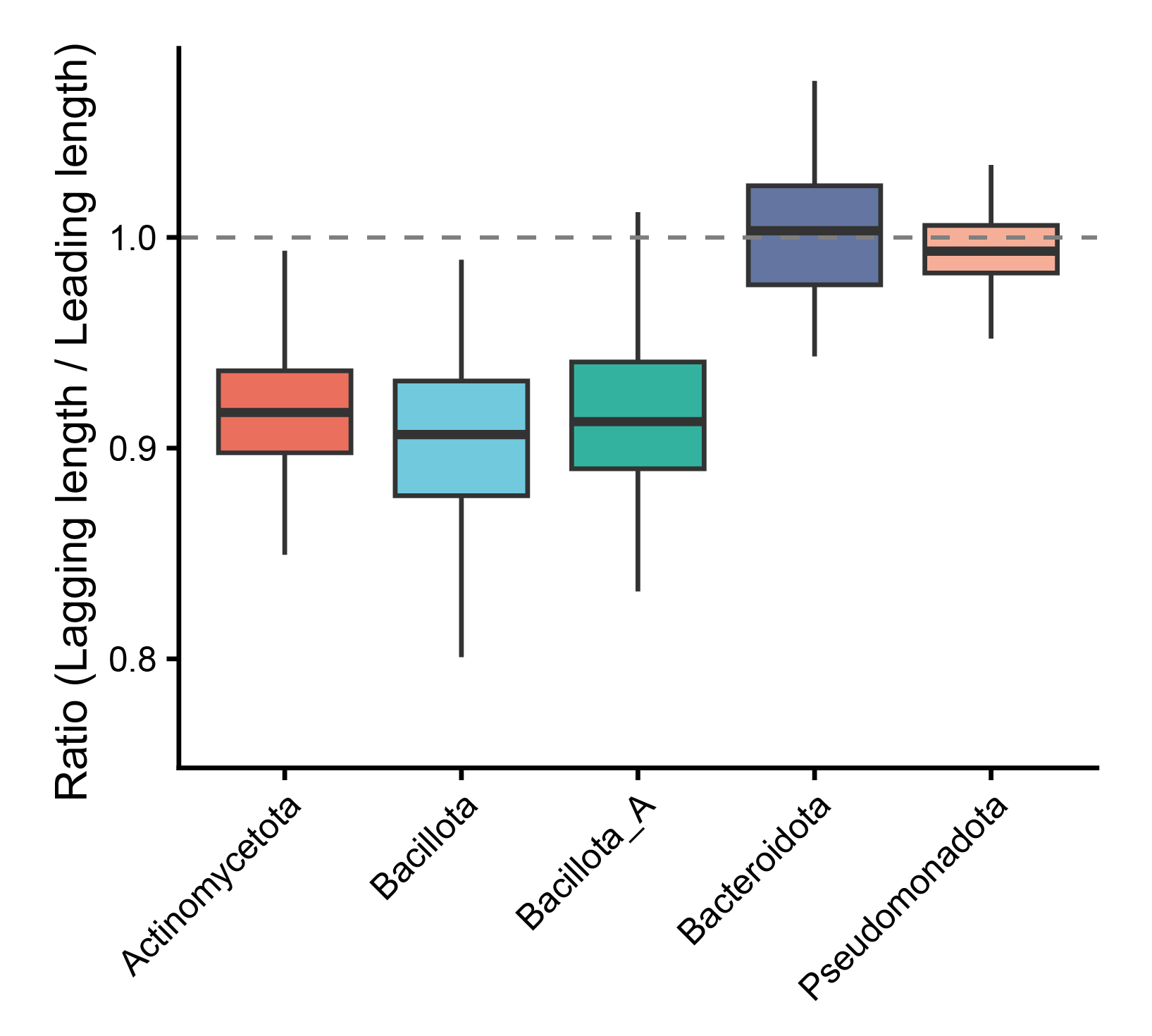

**Fig S4. Strand-specific length asymmetry of genes.** Distribution of the gene length ratio between the lagging and leading strands across the major bacterial phyla. The ratio is calculated for each genome as the average length of genes located on the lagging strand divided by the average length of genes located on the leading strand. The dashed line at 1.0 represents the baseline where average gene lengths are equal on both strands.

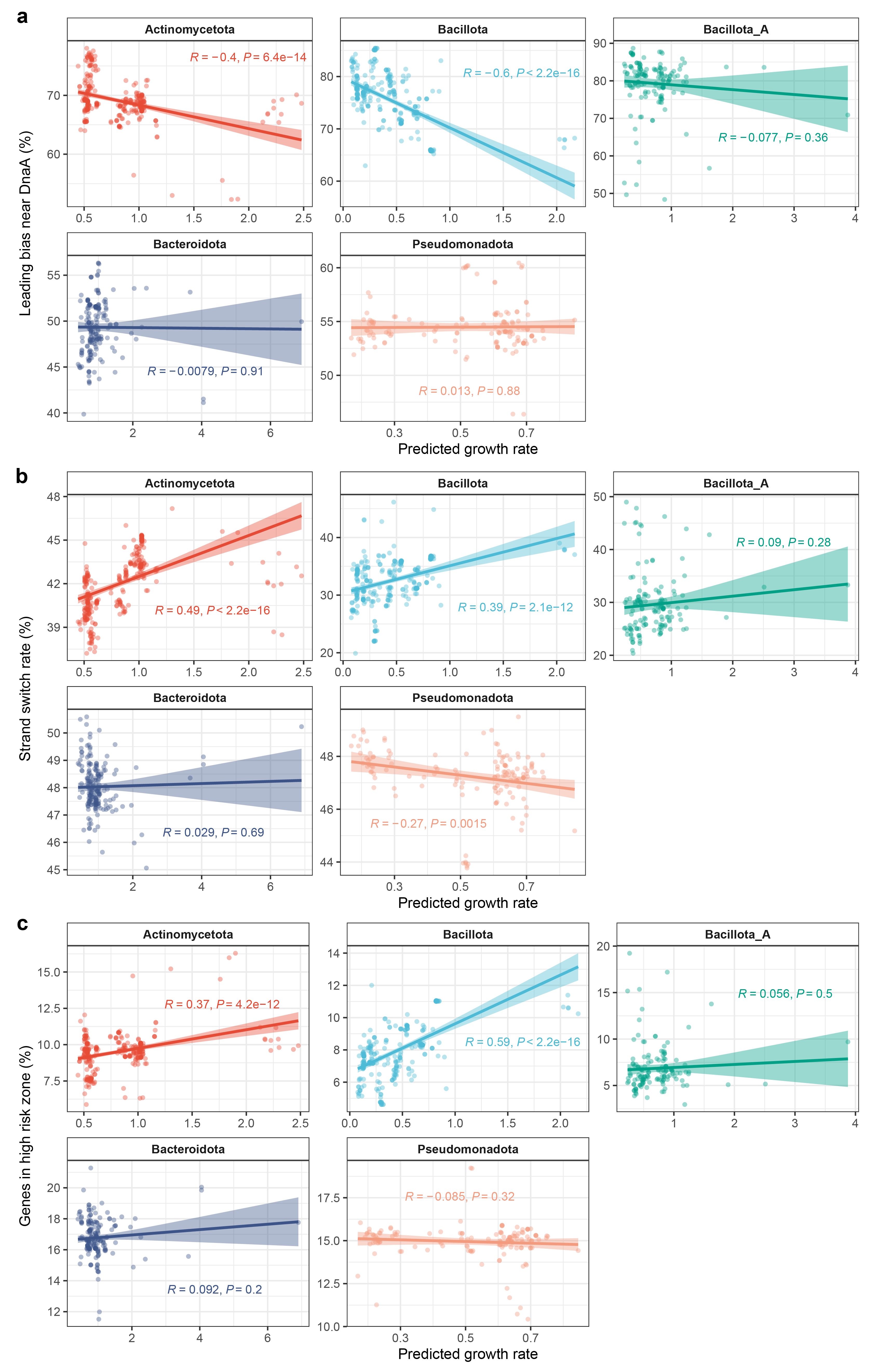

**Fig S5. Relationships between genomic architecture and bacterial growth rate.** Correlations between predicted growth rate and the leading strand bias near *dnaA* (a), strand switch rate (b), and genes in high-risk zone (c) across the major bacterial phyla.

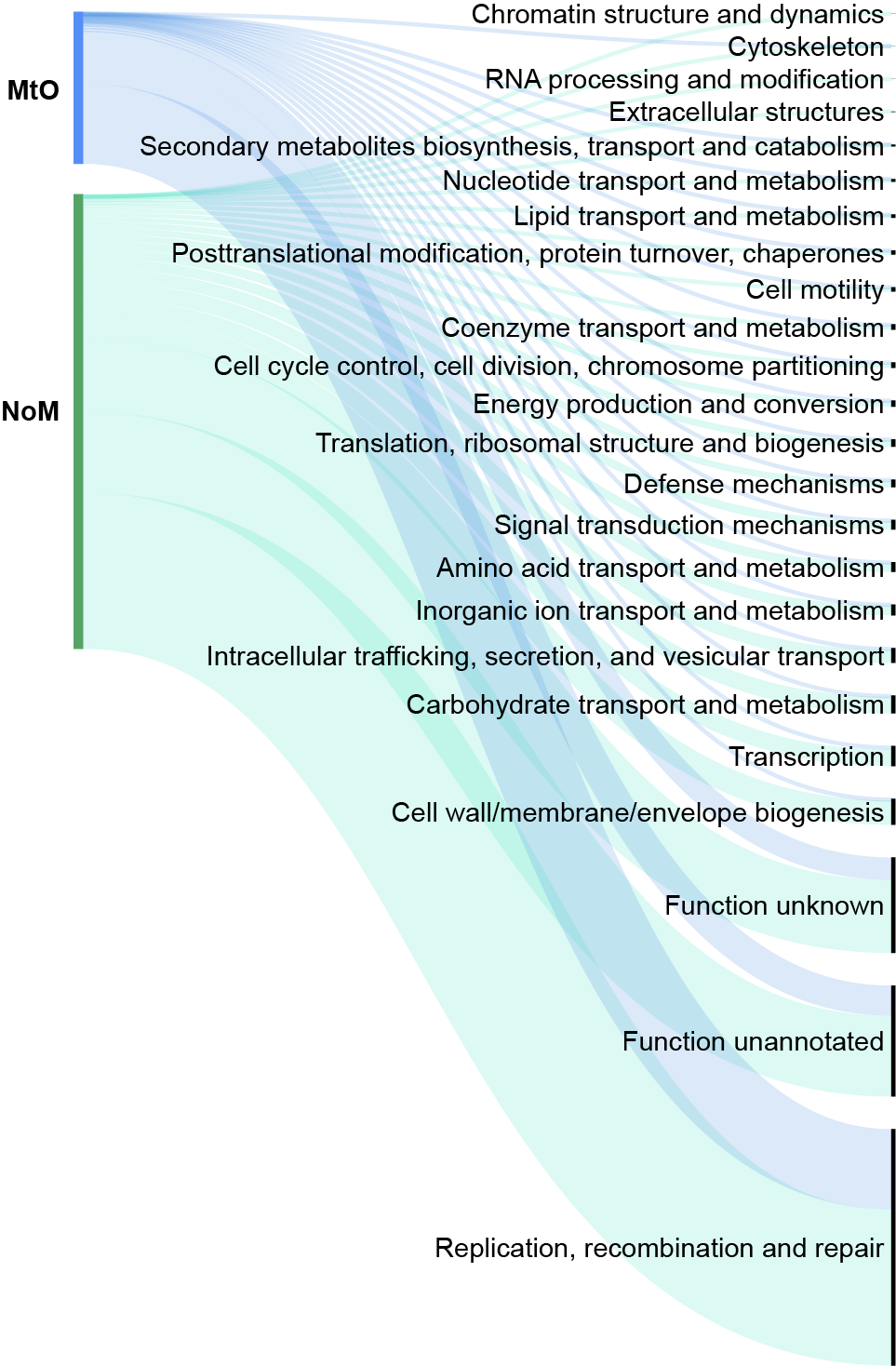

**Fig S6. Distribution of COG function composition of MtO genes and NoM CDSs.** MtO CDSs refer to multi copy CDSs in complete genomes but only one copy in corresponding draft genomes**.** The NoM CDSs refer to CDSs that exist in the complete genomes but have no corresponding match in the draft genomes.

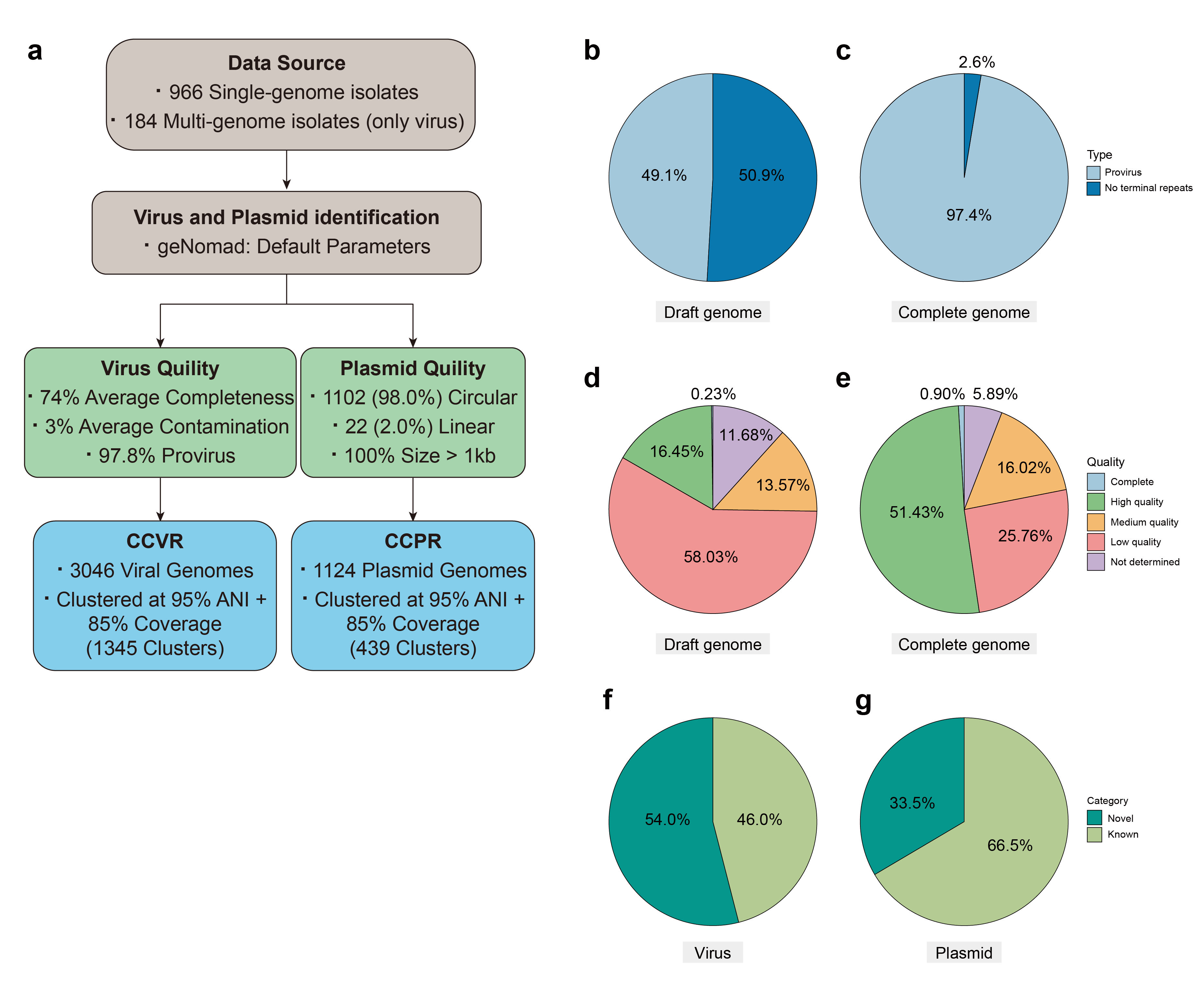

**Fig S7. Overview of phage and plasmid databases.** **a** Flowchart and parameters for phage and plasmid identification. **b** Phage species from draft genomes. **c** Phage species from complete genomes. **d** The quality of CheckV of phages from draft genomes. **e** The quality of CheckV of phages from complete genomes. **f** The percentage of novel phages compared with the database. **g** The percentage of novel plasmids compared with the database.

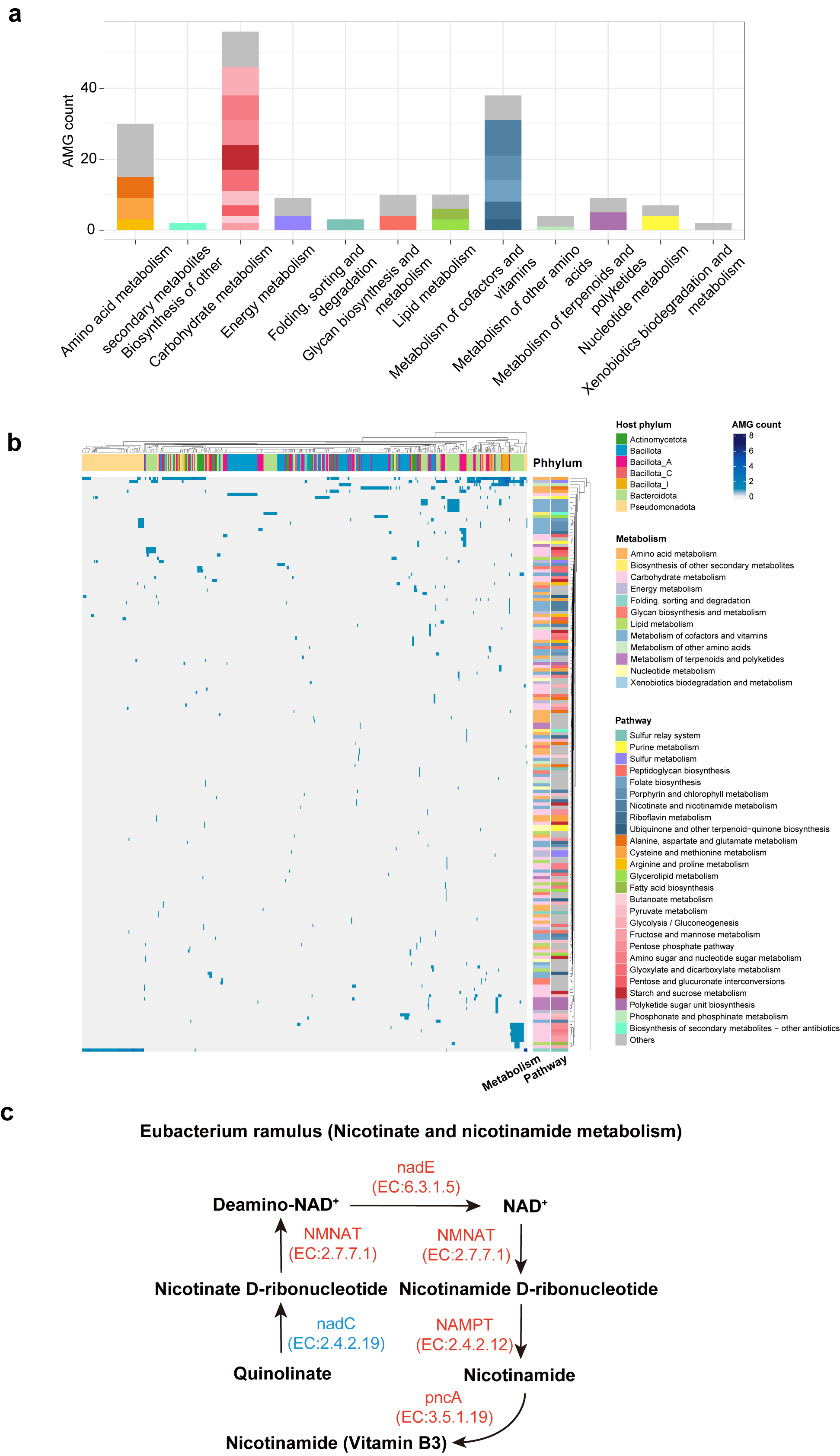

**Fig S8. The functional repertoire of phage-encoded auxiliary metabolic genes (AMGs). a** Distribution of AMG functional categories, dominated by cofactor/vitamin metabolism and carbohydrate metabolism. **b** Heatmap profiling of the presence of AMGs across different host phyla. **c** A schematic model of the nicotinate and nicotinamide metabolism pathway in *Eubacterium ramulus*, where phage-encoded enzymes (red) complement host-encoded enzymes (blue) to facilitate NAD^+^ biosynthesis, exemplifying a synergistic phage-host metabolic interaction.

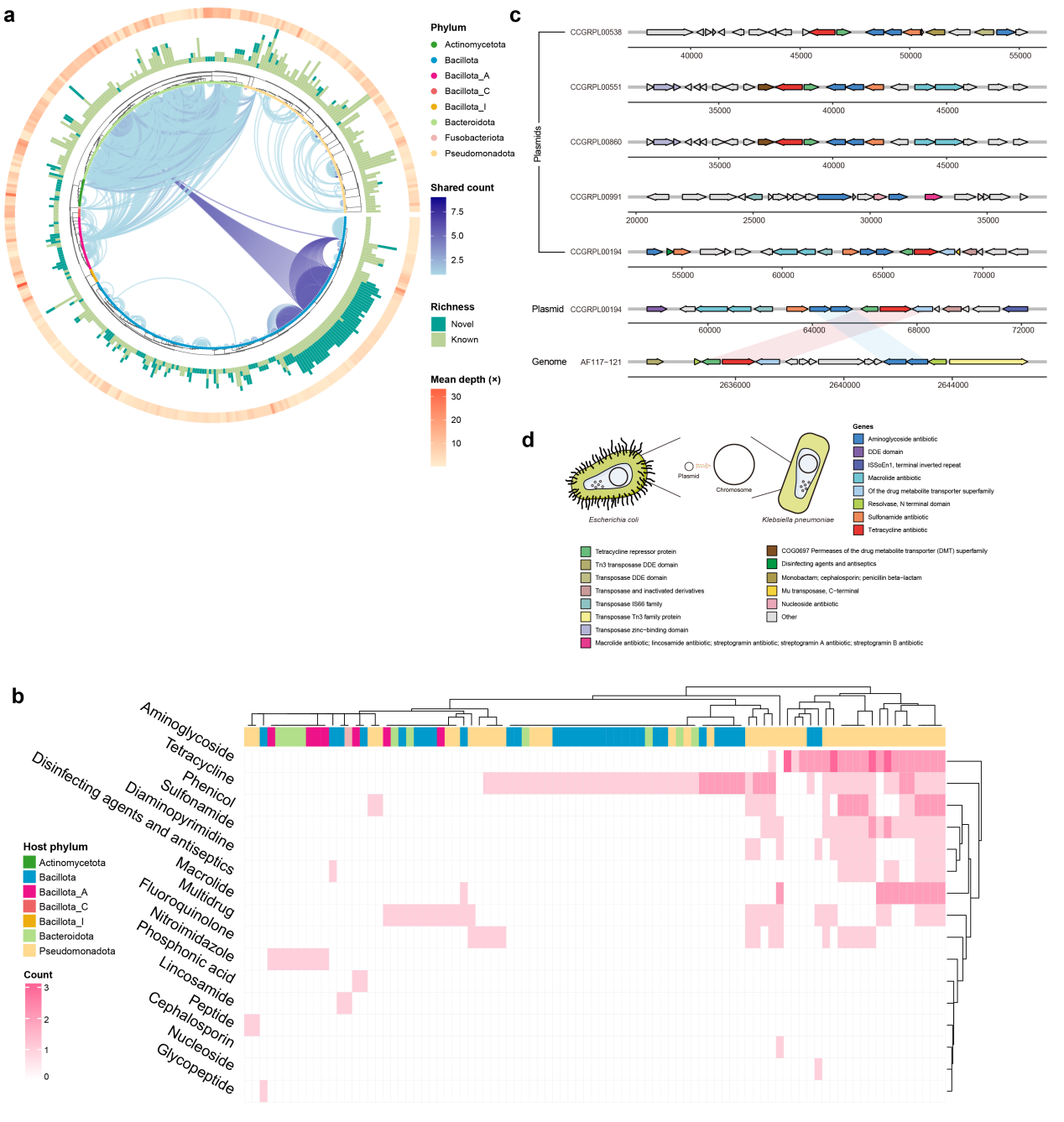

**Fig S9. Cross-taxa dissemination of plasmids facilitates the spread of antibiotic resistance genes. a** The transmission range of plasmids was plotted based on phylogenetic tree of CCGR. The blue lines in the inner layer indicate that there are plasmids from the same cluster in different bacterial genomes. The middle layer shows the richness of the displayed plasmids, that is, the number of plasmids clusters. Dark green indicates novel plasmids in our database, and light green indicates previously described plasmids. The outermost layer shows the copy numbers of the plasmid. **b** Heatmap profiling the distribution of antibiotic resistance genes (ARGs) across different host phyla. **c** Clustered transfer of resistance genes present on plasmids. **d** An example of resistance gene transfer from a plasmid of *Escherichia coli* to *Klebsiella pneumoniae*.

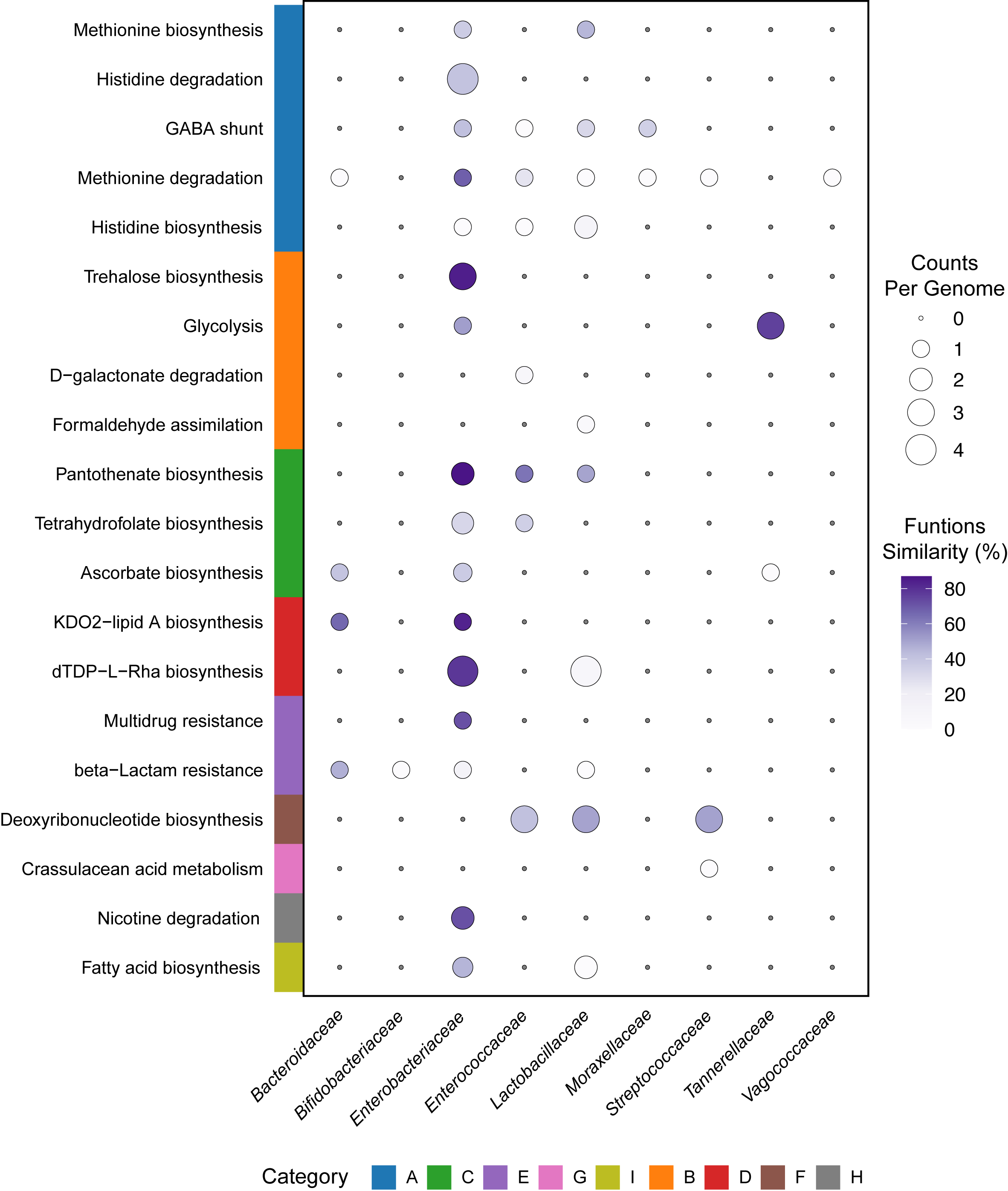

**Fig S10. Enrichment of plasmid-carried functional modules across representative gut bacterial families.** Bubble size corresponds to the average gene count per bacterial genome, while color intensity represents the functional similarity score (%). Functional categories are indicated by the colored bar on the left.

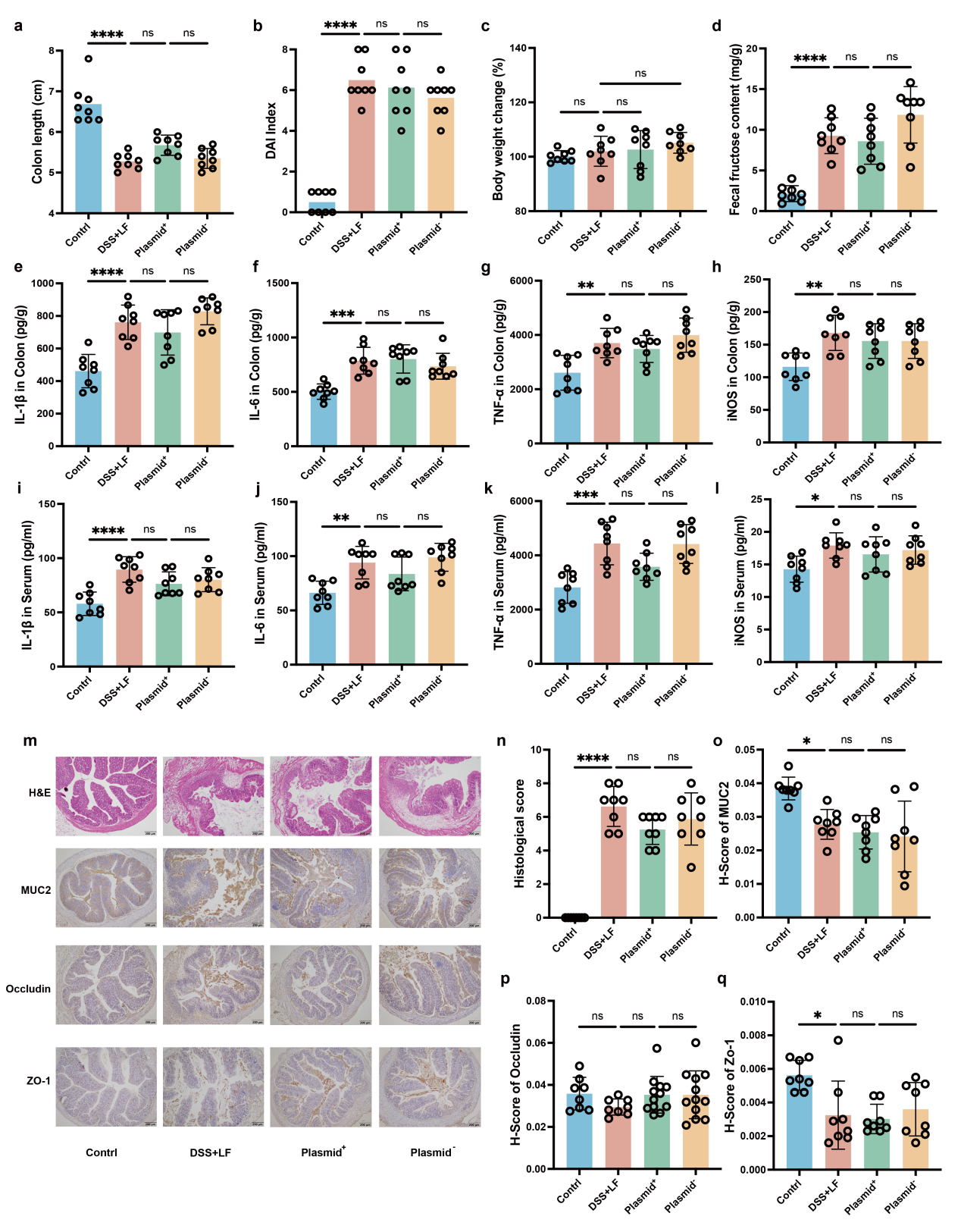

**Fig S11. The efficacy of Plasmid^+^ *L. brevis* is abrogated under low-fructose conditions.** Assessment of colitis severity in mice fed a low-fructose (LF, 1%) diet and challenged with DSS. Groups included Control, DSS+LF, and DSS+LF treated with either Plasmid^+^ or Plasmid^-^ *L. brevis*. **a-c** Macroscopic markers of colitis: colon length (a), DAI scores (b), and body weight change (c). **d** Fecal fructose content, showing no significant accumulation or depletion across groups due to low dietary input. **e-l** Inflammatory cytokine profiles in colonic tissue (e-h) and serum (i-l). **m** Representative histological images of H&E staining and IHC for MUC2, Occludin, and ZO-1. Scale bars, 200 µm. **n-q** Quantitative histological scoring (n) and optical density analysis of barrier proteins (o-q). Data are presented as mean ± s.e.m. Statistical significance was determined by one-way ANOVA with Tukey’s post hoc test. **P* < 0.05, ***P* < 0.01, ****P* < 0.001, *****P* < 0.0001; ns, not significant.

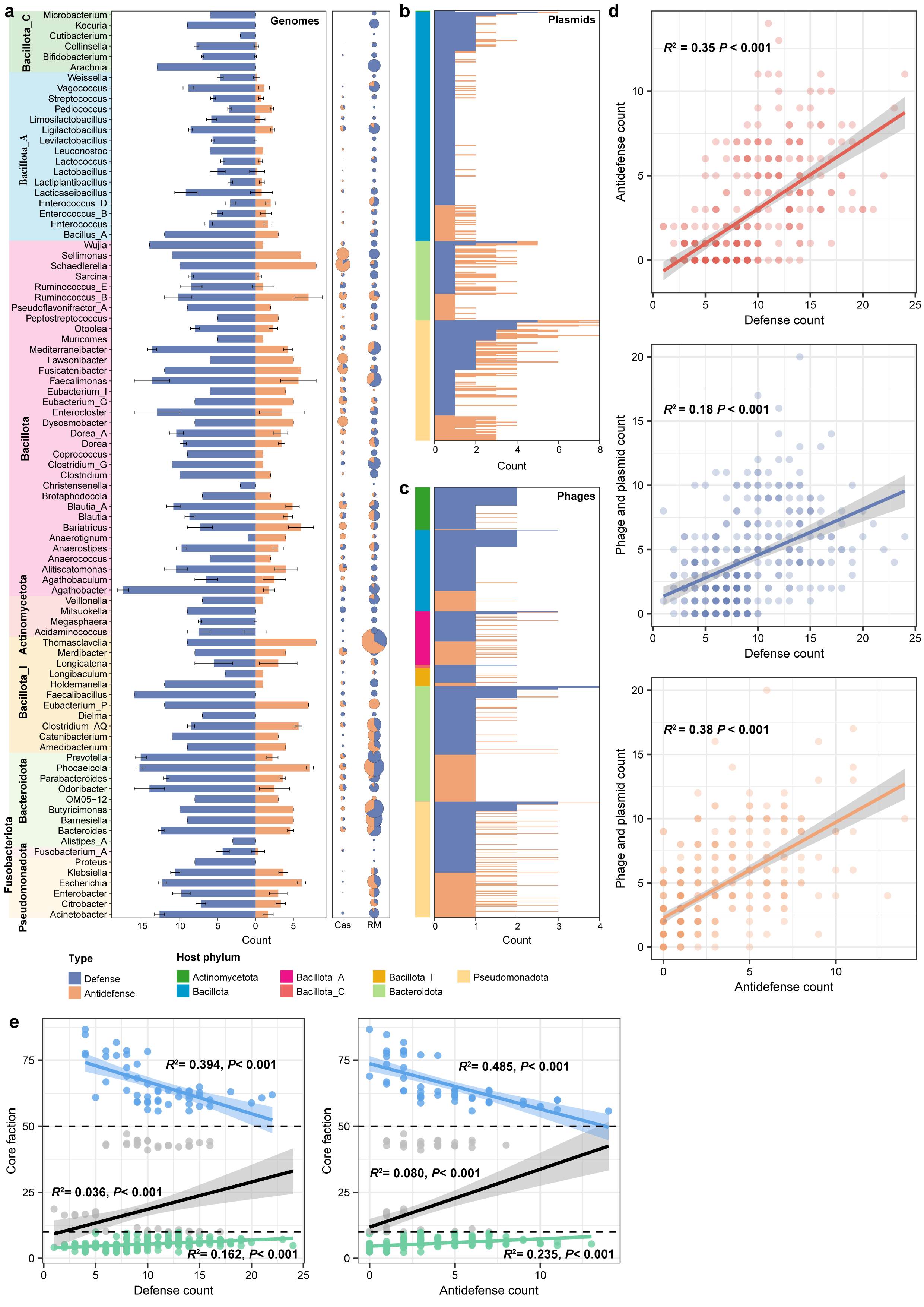

**Fig S12. Complete genome reveals the relationship between defense systems and evolution.** **a** Distribution of defense systems and anti-defense systems in different genera; the proportion of the two types of defense systems, Cas and RM, is shown. **b** Distribution of defense and anti-defense systems on plasmid sequences. **c** Distribution of defense and anti-defense systems on phage sequences. **d** Linear regression analysis of defense systems and mobile genetic elements. **e** Correlation analysis between defense systems or anti-defense systems and core gene fractions.

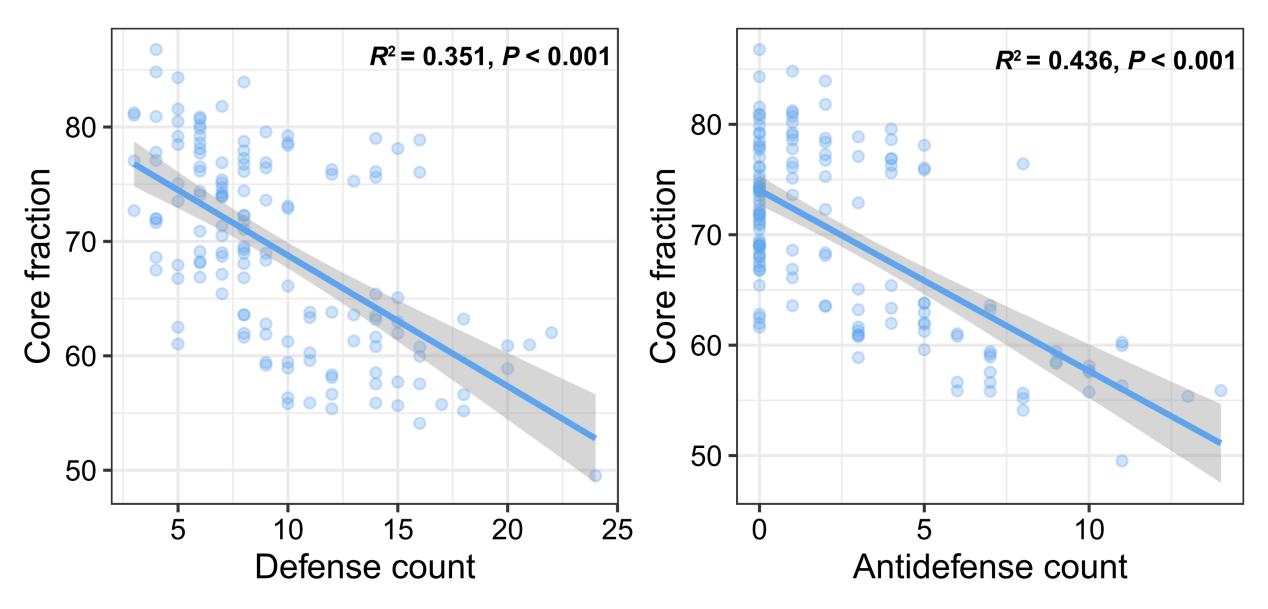

**Fig S13. Relationships between core genes and defense systems/anti-defense systems.** Linear correlation at the species level.

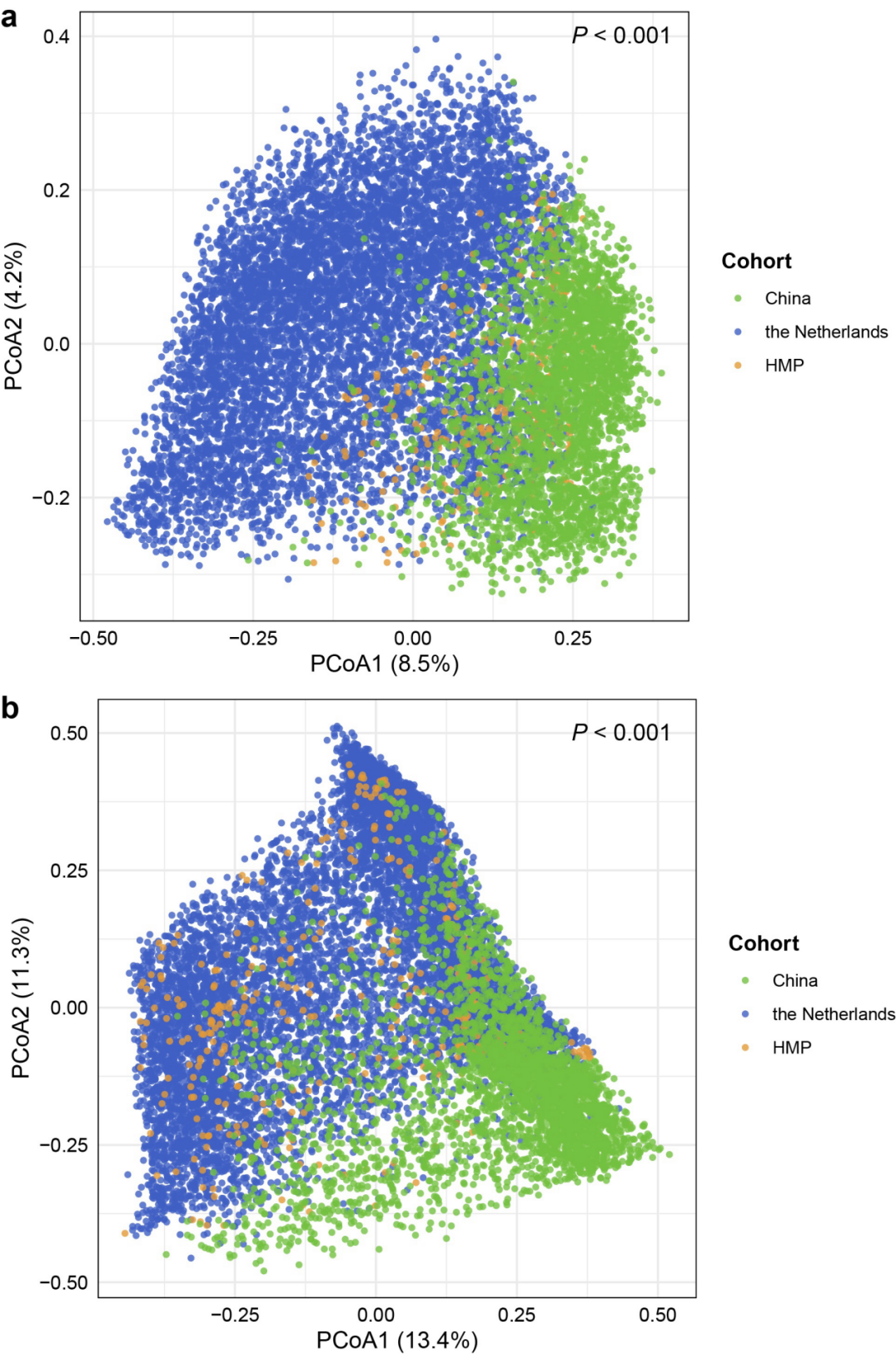

**Fig S14. Composition of phages and plasmids in three healthy cohorts.** Variations in phage (a) and plasmid (b) composition based on the Bray-Curtis dissimilarity between cohorts from China, the Netherlands, and HMP shown in PCoA plots and tested by PERMANOVA.

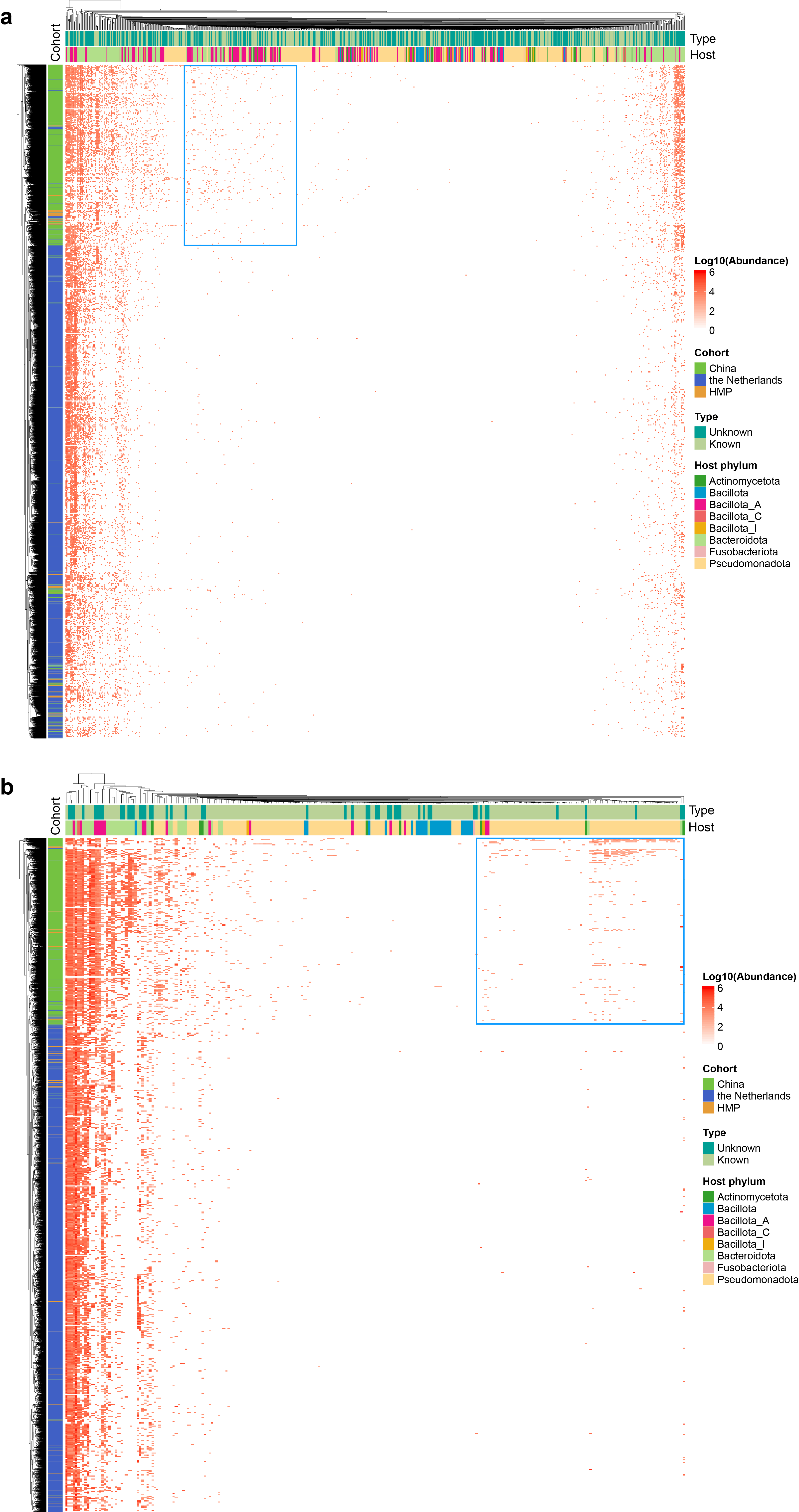

**Fig S15. Abundance distribution of phages and plasmids in three cohorts.** Heatmaps showing the differences in the abundance of phages and plasmids between cohorts from China, the Netherlands, and HMP.

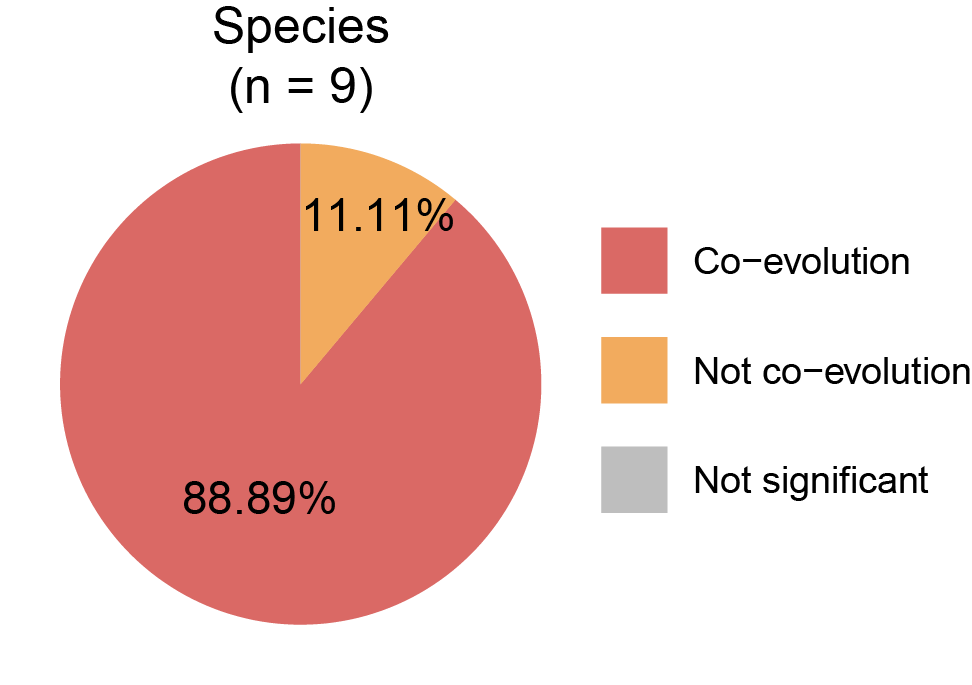

**Fig S16. Co-evolution between host genomes, phages, and plasmids.** Pie chart showing the proportions of co-evolution, not co-evolution and not significant cases among nine species.
