## Supplementary material for "Complete Genomes of Cultivated Gut Bacteria Reveal Mobile Genetic Element-Driven Functional Diversity with Therapeutic Implications": STables: STable info.docx

**Description of Additional Supplementary Files**

**File Name:** Supplementary Data 1

**Description:** Statistics for sequencing data of the 1,150 complete genomes.

**File Name:** Supplementary Data 2

**Description:** Taxonomic information of the 1,150 complete genomes.

**File Name:** Supplementary Data 3

**Description:** Comparison of genomic metrics for 966 complete genomes and paired draft genomes.

**File Name:** Supplementary Data 4

**Description:** Comparison of protein function differences between the draft genome and the finished genome.

**File Name:** Supplementary Data 5

**Description:** The information of phages.

**File Name:** Supplementary Data 6

**Description:** The information of plasmids.

**File Name:** Supplementary Data 7

**Description:** Types and quantities of defense systems of strains, plasmids, and phages.

**File Name:** Supplementary Data 8

**Description:** Composition of mice diet.

**File Name:** Supplementary Data 9

**Description:** Scoring systems for disease activity index (DAI).

**File Name:** Supplementary Data 10

**Description:** Scoring systems for histological score.

**File Name:** Supplementary Data 11

**Description:** The cohorts used of this study.
